# Post translational modification of duplicated ribosomal protein paralogs promotes alternative translation and drug resistance

**DOI:** 10.1101/2021.10.06.463374

**Authors:** Mustafa Malik-Ghulam, Mathieu Catala, Michelle S. Scott, Sherif Abou Elela

**Affiliations:** Département de microbiologie et d’infectiologie, Faculté de médecine et des sciences de la santé, Université de Sherbrooke, Sherbrooke, QC J1E 4K8, Canada; Département de biochimie, Faculté de médecine et des sciences de la santé, Université de Sherbrooke, Sherbrooke, QC J1E 4K8, Canada

## Abstract

Ribosomes are often seen as monolithic machines produced from uniformly regulated genes. However, in yeast most ribosomal proteins are produced from duplicated genes. Here, we demonstrate that gene duplications may serve as a stress response mechanism that modulates the global proteome through differential post-translational modification of ribosomal proteins paralogs. Our data indicate that the yeast paralog pair of the ribosomal protein L7/uL30 produces two differentially acetylated proteins. Under normal conditions most ribosomes incorporate the hypo-acetylated ‘major’ form favoring the translation of genes with short open reading frames. Exposure to drugs, on the other hand, increases the production of ribosomes carrying the hyper-acetylated minor paralog that increases translation of long reading frames. Many of these genes encode cell wall proteins that increase drug resistance in a programed change in translation equilibrium. Together the data reveal a mechanism of translation control through the differential fates of near-identical ribosomal protein isoforms.

## Main

Ribosomes are ribonucleoprotein complexes required for protein synthesis ^1, 2^. The basic structure of the ribosome is conserved from bacteria to human with increasing complexity in terms of the number and sizes of rRNAs and proteins in higher eukaryotes ^2–5^. Ribosomes are viewed as universal machines built for precision and mass production ^6–9^. However, this view is being challenged by observations of variations between ribosomal protein gene (RPGs) in terms of their expression, regulatory pathways, and variable incorporation into ribosomes ^10–17^. While the concept of specialized ribosome is debated, the heterogeneity of ribosome composition and regulatory programs is irrefutable. Most eukaryotes have variable amounts of mRNA produced by the different RPGs and mass-spectrometer analyses continues to detect variations in the ribosome populations extracted from different tissues and growth conditions ^10, 18^.

In the yeast *S. cerevisiae*, ribosomal protein genes (RPGs) are mostly produced from independently regulated duplicated genes (Extended Data Fig. 1)^18^. Out of the 137 yeast RPGs, 118 (56 pairs) are duplicated and 101 have introns that must be removed to produce the mRNA ^18^. Paralogous ribosomal protein pairs are generally more than 95% identical. However, deleting individual paralogs give different phenotypes, suggesting functional specialization. The origin of this proposed paralog specialization remains unclear and actively debated ^19–21^.

Here we systematically assess *S. cerevisiae* paralog functions with respect to their expression levels and amino acid differences. For the most disparate pairs, in terms of expression or sequence, we used gene conversion to create simplified ‘homogenized’ strains, in which each individual duplicated RPG was expressed both from its natural locus and as a replacement for the similar paralog and tested them for growth under different conditions. Overall, our results indicate that duplicated ribosomal protein genes (dRPGs) regulate drug resistance through paralog-specific effects on translation.

## Results

### Gene conversion of duplicated ribosomal protein genes reveal paralog subfunctionalization

Comparing the expression level of the 29 RPG pairs with significant difference in both regulatory and coding sequence using qRT-PCR, we identified four 60S and one 40S RPG pairs responding differently to changes in growth conditions (Fig. 1a and Extended Data Fig. 1a-b). Most differences between the gene pairs are found in the regulatory sequence (< 70% homology) controlling their expression (Extended Data Fig. 1b). Consistently, most gene pairs produced different amounts of RNA and proteins (Fig. 1b-c). In general, paralogs producing most RNA also produced most protein except in the case of *eL37/RPL37* paralog, which are differentially translated ^22^. In all cases, one of the two paralogous proteins (the major form) dominated over the other (the minor form) by 3 to 4 times, suggesting that most of the ribosomes are produced from one gene copy (Fig. 1c). Deleting one of the two dRPGs copies reduced the abundance of the mRNA generated by the pair, except in the case of *uL30/RPL7* and the major copy of *eL36/RPL36* (Fig. 1d). Homogenization of dRPGs, except *el27bb/rpl27bb*, did not decrease mRNA coding for their associated proteins (AA and BB, Fig. 1d). We conclude that the expression level of RPGs is not strictly dependent on gene duplication or the paralog identity.

**Figure 1.**
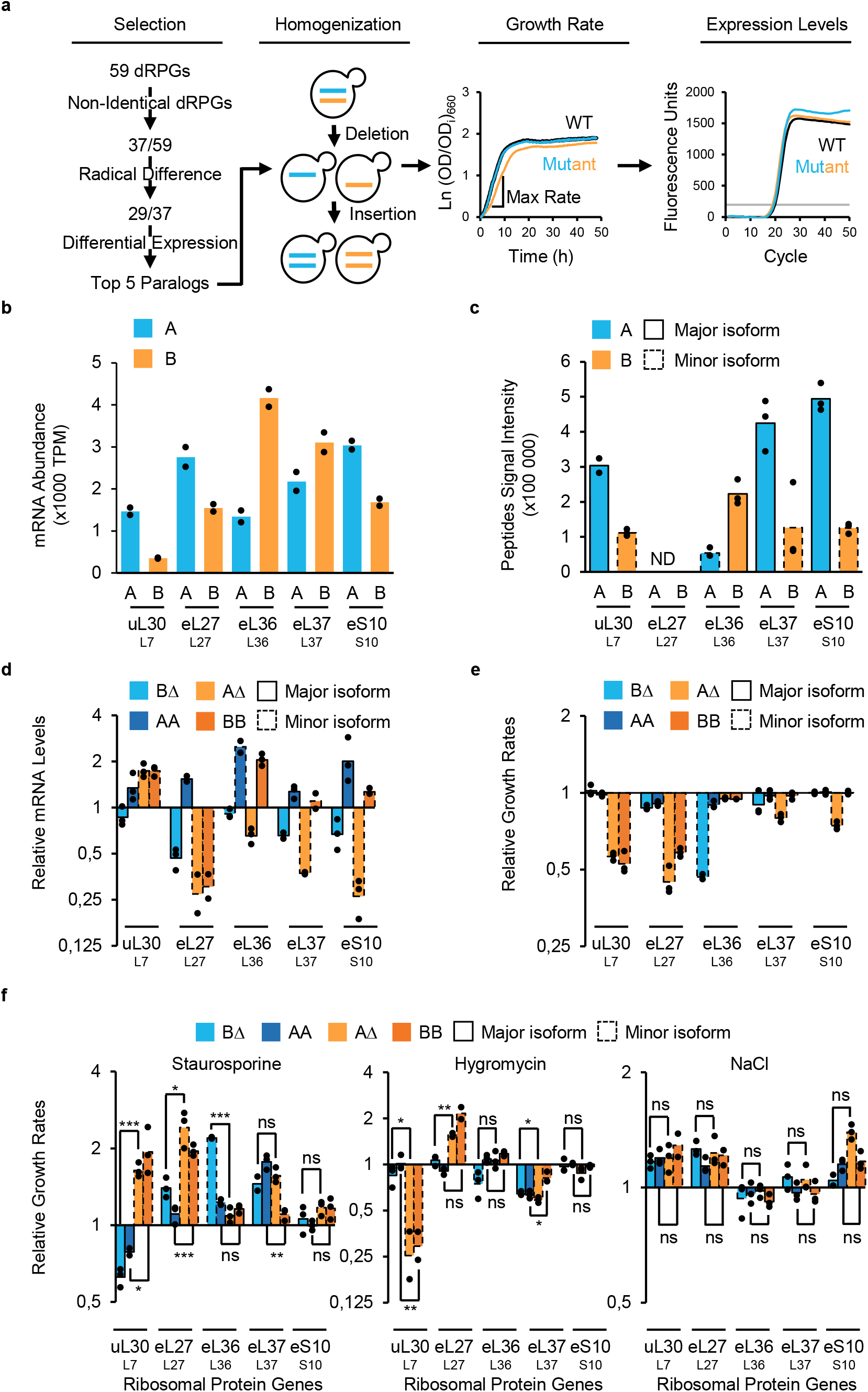
Homogenization of yeast ribosomal proteins identifies dose independent paralogs subfunctionalization. **a,** Gene conversion strategy for “homogenization” of ribosomal protein genes. Genes showing paralog-specific response to different conditions were selected for homogenization. Homogenized strains were created by replacing one paralog locus including all regulatory sequences, introns and UTRs, with sequence of the other. Cells carrying one or two copies of a single paralog were examined for growth in liquid media and maximum growth rates were calculated for each strain. Duplicated ribosomal protein genes (dRPG) expression level was examined by qRT-PCR to identify dose-dependency of growth defects. **b,** RNA abundance in transcripts per million (TPM) of each paralog obtained by RNA sequencing for 2 biological repeats. Universal and conventional names of each protein are shown below. **c,** Abundance of protein isoforms generated by each dRPG was determined by Swath Multiple Reaction Monitoring (MRM) identifying the major and minor isoforms for 3 biological repeats. No suitable peptides that could reliably distinguish between L27 paralogs were found, hence their peptide signal intensities were not determined (ND). **d,** Total RNA generated by each gene pair was detected using primers common to both paralogs in qRT-PCR after deletion or homogenization of dRPGs and is shown relative to mRNA detected in wild-type strain. BΔ and AΔ indicate RNA detected in strains lacking the B or A paralog, while AA and BB indicate RNA detected in cells containing two copies of the A or B isoforms respectively. **e,** Growth rates of deletion and homogenized strains were determined and shown relative to wild-type. **f,** Growth rates of deletion and homogenized strains were determined in media containing staurosporine (3 µg/ml), hygromycin (100 µg/ml) or NaCl (0.9 M) and the effect on growth are shown relative to wild type treated similarly. Bars shown in **c**-**f** represent the means of 3 biological replicates shown as data points (*p<0.05, **p<0.01, ***p<0.001 by two-tailed unpaired t-test assuming unequal variance).

As expected, the deletion of the major paralog, which produces most RPs in the cell, impaired cell growth in rich media (Fig. 1e). This growth defect is restored by the expression of two copies of the minor paralog except in the case of *uL30* and *eL27/RP27*, where the minor paralog duplication fails to complement the deletion of the major copy (Fig. 1e). Exposure to stress revealed different set of copy specific effects that could not be explained by reduced gene expression or growth rate. For example, deletion of *eL27/RPL27* major paralog, which reduces expression and inhibits growth under normal conditions, enhanced growth in the presence of staurosporine and hygromycin (Fig. 1f). In contrast, the minor paralog of *uL30*, which does not inhibit gene expression or cell growth, was both necessary and sufficient for cell resistance to staurosporine (Fig. 1f and Extended Data Fig. 2a). These data indicate that dRPGs are only partially redundant and identify *uL30* as a good model for studying the functional specialization of dRPGs.

### *uL30* modulates cell growth and ribosome biogenesis in a paralog-dependent manner

The proteins generated from *uL30* dRPGs differ by five amino acids, 4 of which are clustered in the N-terminus, leading to major difference in the predicted secondary structure of the N-terminal domain (Extended Data Fig. 1b and 2b). The protein featuring the more structured N-terminal domain (uL30A) is more abundant forming the bulk of the ribosome in the cell making it more of the housekeeping version of the pair (Extended Data Fig. 2d). Consistently, the expression of this major paralog (*uL30A*) is required for normal growth and assembly of the 60S subunit, while the minor paralog (*uL30b*) is only required for drug resistance (Fig. 2a-d). Cells expressing one or two copies of the A version (*ul30bΔ* and *ul30aa, respectively*) were more sensitive to staurosporine and produced shorter polyribosome than wild type-cells but otherwise were normal (Fig. 2a-d and Extended Data Fig. 3). In contrast, cells expressing the B version (*ul30aΔ* and *ul30bb*) were more resistant than the wild type to staurosporine but grew slower under normal growth conditions. This caused a subunit imbalance and an accumulation of 40S subunit awaiting the 60S subunit, or ‘half-mers’ (Fig. 2a-c and Extended Data Fig. 3).

**Figure 2.**
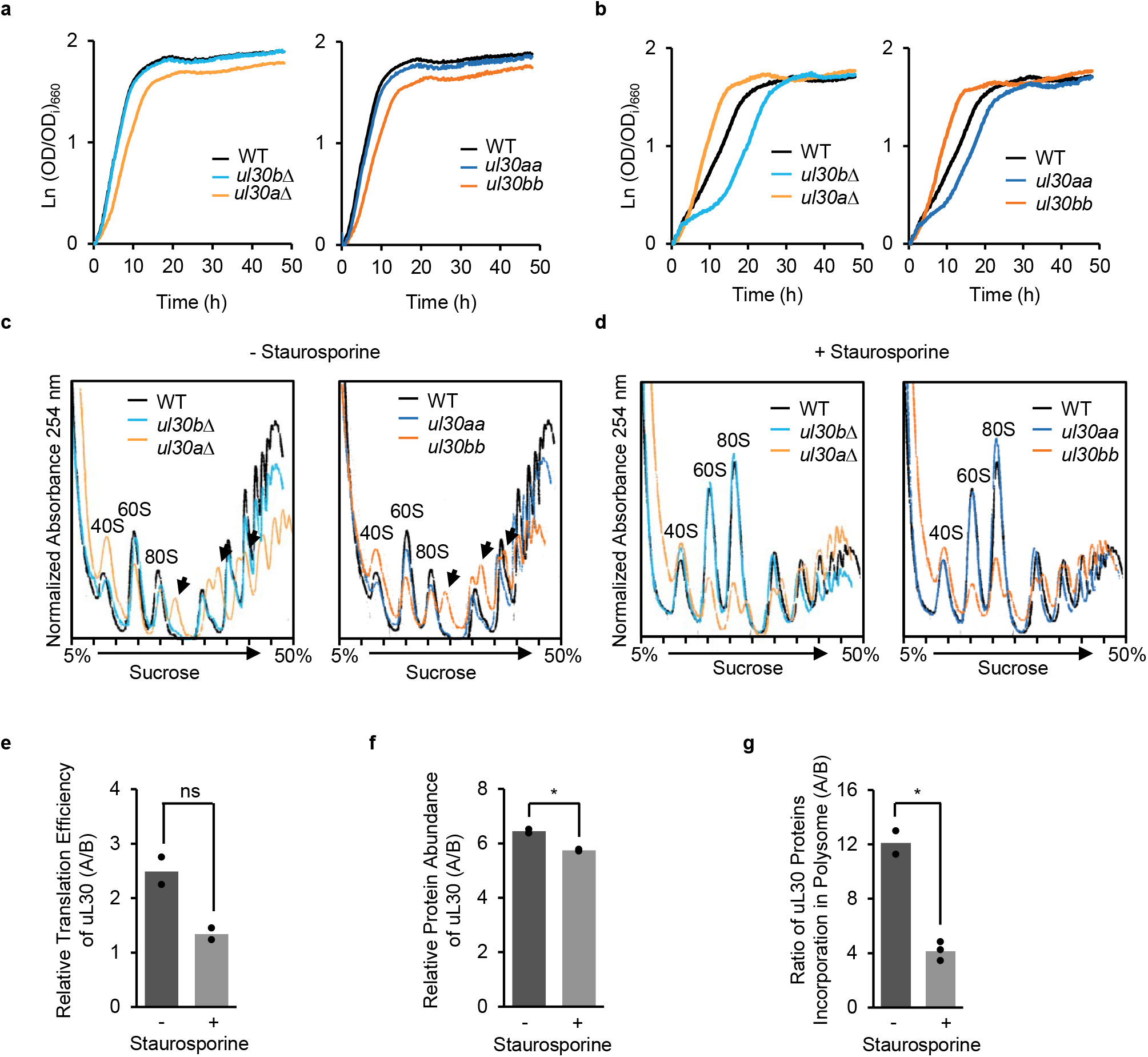
*uL30* paralogs differentially alter cell growth, ribosome biogenesis and staurosporine resistance. **a,b,** Growth curves of wild-type (WT), deletion strains (*ul30bΔ* and *ul30aΔ*) and homogenized strains (*ul30aa* and *ul30bb*) grown in complete synthetic media without (**a**) or in the presence of staurosporine (**b**). **c,d,** Polysome profiles were obtained from WT, cells lacking one paralog or expressing two copies of same paralog from cultures grown in complete synthetic media without (**c**) or in the presence of staurosporine (**d**). Position of 40S and 60S ribosomal subunits and 80S ribosomes are indicated. Arrows indicate position of half-mers or 40S awaiting the 60S subunit. Curves shown in **a**-**d** are representative examples from 3 biological repeats. **e**, Translation index (mRNA associated with polyribosomes / mRNA associated with monosome and subunits) was determined using digital droplet PCR for each *uL30* paralog in absence (-) or presence (+) of staurosporine and the ratio for uL30A over uL30B is reported for 2 biological repeats (ns: p=0.078 by paired two-tailed t-test). **f**, Protein abundance of uL30 was determined by MRM in absence or presence of staurosporine and the ratio of uL30A over uL30B is reported for 2 biological repeats (*p <0.05 by paired two-tailed t-test). **g**, The amount of uL30 proteins incorporated into ribosomes was determined using MRM in absence (n=2) or presence (n=3) of staurosporine and the ratio of uL30A over uL30B is reported (*p <0.05 by two-tailed t-test with unequal variance).

Surprisingly, exposure to staurosporine inhibited translation in a paralog-dependent manner. Staurosporine significantly reduced the size of the polyribosome in cells expressing uL30A but had no effect on those expressing uL30B (Fig. 2d and Extended Data Fig. 3b). The paralog-specific phenotypes including the response to staurosporine are not linked to the depletion of ribosomal proteins since most strains expressed uL30 and other ribosomal proteins to similar levels (Extended Data Fig. 2c and g). Furthermore, the combination or choice of uL30 paralog did not significantly alter the expression of most RPGs arguing that the subunit imbalance detected in the absence of the A form is due to defects in the 60S assembly, which is consistent with the role of uL30 in ribosome biogenesis. Exposing cells to staurosporine altered the ratio of *uL30* proteins by favouring the translation of the minor paralog and its incorporation into active ribosomes (Fig. 2e-g). Preferential translation of *uL30A* under normal conditions could be explained by its higher Kozak score and the clustering of suboptimal codons at the 5’ end of *uL30B* (Extended Data Fig. 4). We conclude that *uL30*A is required for the biogenesis of the 60S subunit, while uL30B is mostly needed for cell resistance to staurosporine.

### The paralogs of *uL30* differentially modulate the translation of defined subsets of mRNA

Comparing the translation pattern of cells expressing two copies of *uL30A or uL30B* identified a set of paralog dependent genes. The ribosome associated mRNAs, in wild type, *ul30aa* and *ul30bb*, was sequenced before or after staurosporine treatment and the results confirmed using RT-qPCR as indicated in Fig. 3a and Extended Data Fig. 5f. Wild-type and *ul30aa* cells displayed similar translation pattern with only 96 genes more- or less-translated in *ul30aa* cells (Fig. 3b-c). In contrast, 2094 genes were more-translated and 458 less-translated in *ul30bb* when compared with wild-type cells (Fig. 3b-c). Strikingly, a set of 16 genes were inversely regulated by *uL30* paralog underlining the paralog specific effect on translation (Supplementary Table 3). The paralog identity had little effect on RNA abundance and the changes in translation did not directionally correlate with changes RNA abundance (Extended Data Fig. 5a, b and d). Therefore, the paralog-specific phenotype is mostly related to changes in translation. The increase in translation index correlated well with the increase in protein amount confirming that the increased association of mRNA with the uL30B containing ribosome lead to increased protein production (Extended Data Fig. 5e). Together these data indicate that uL30 alters translation in a paralog-dependent manner.

**Figure 3.**
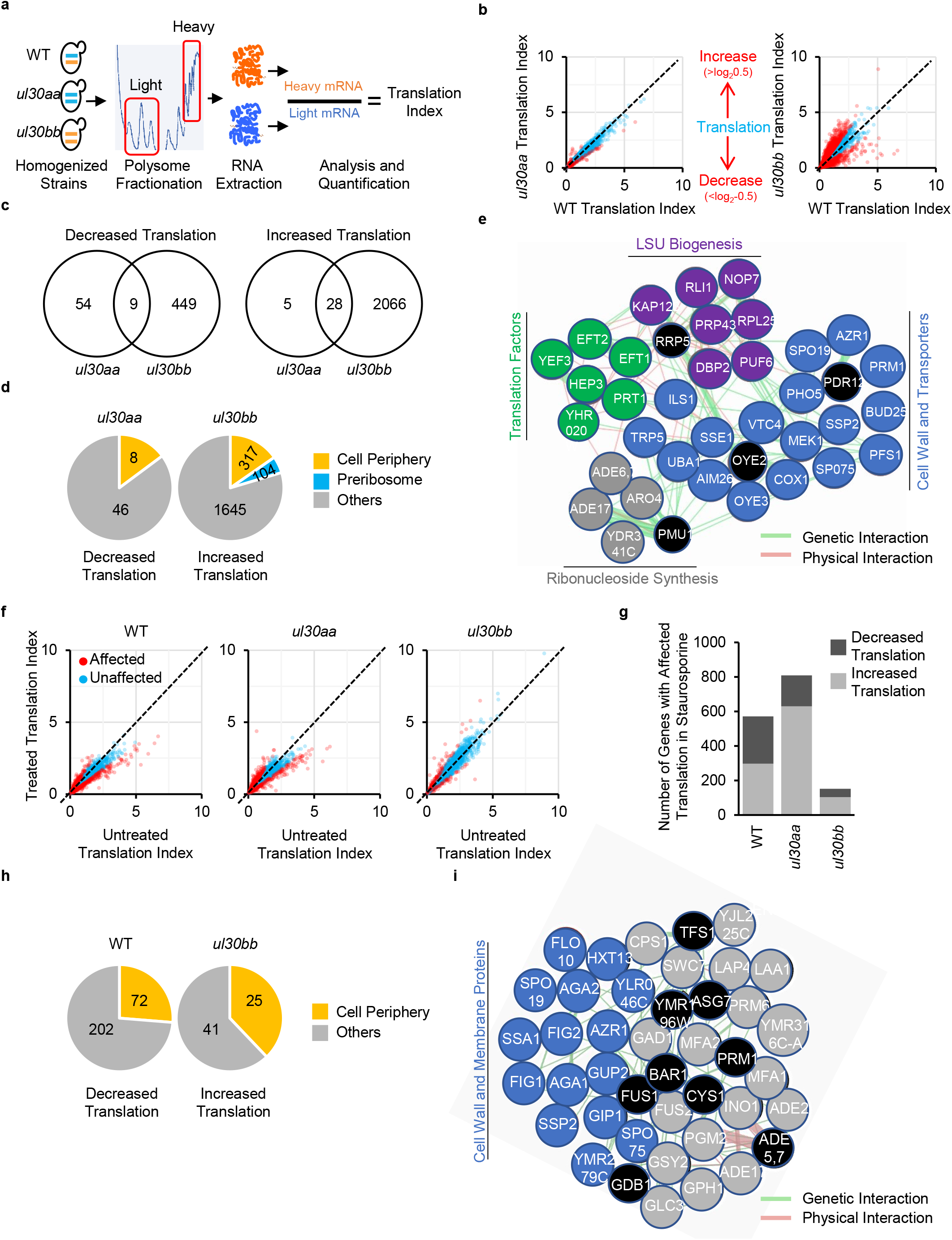
Exposure to staurosporine modulates the translation of genes coding for mitochondrial and cell periphery genes in a paralog dependent manner. **a,** Strategy for determining translation profiles of homogenized *uL30* strains. mRNA was extracted from heavy (polyribosome) and light (monosome and subunit) fractions and sequenced. Translation index was calculated as the ratio of mRNA associated with heavy to light fractions. **b,** Translation index of mRNAs (expression >1 TPM) in homogenized strains is compared to that of wild type. mRNAs showing differential translation by more than log_2_ 0.5 difference are shown in red. **c,** Venn diagram of number of genes with changed association to polysomes comparing homogenized strains (*ul30aa* and *ul30bb*) to WT identified in (**b)**. **d,** Distribution of the number of genes present in enriched component gene ontology categories (p <0.001) for genes with specific changes in association to polysomes. **e,** Map of the genetic and physical interactions of the top up-translated genes in *ul30bb* strain was generated with the Genemania network construction tool. Up-translated genes were grouped and colored by functional pathways, genes in black circles are not affected by *ul30bb*. **f,** Translation index of mRNAs (expression >1 TPM) in WT and homogenized strains is compared in cells grown in absence (untreated) or in presence of staurosporine (treated). mRNAs showing differential translation by more than log_2_ 0.5 difference are shown in red. **g,** Bar graph showing the number of genes under-translated (dark grey) or over-translated (light grey) in response to staurosporine. **h,** Distribution of the number of genes present in enriched component gene ontology categories (p <0.001) for genes with specific changes in association to polysomes after staurosporine treatment. **i,** Map of the genetic and physical interactions of the top up-translated genes in *ul30bb* strain in presence of staurosporine was generated with the Genemania network construction tool. Up-translated genes associated to cell wall and membrane component category were grouped and colored in blue, genes in gray associate to other categories and genes in black circles are not affected by *ul30bb*. RNA sequencing has been performed on 2 biological repeats.

### The minor paralog of *uL30* induces the translation of genes associated with cell periphery

Gene ontology analysis identified cell periphery as the only inversely regulated category by *uL30 paralog*. The expression of the A form decreases the translation of cell periphery genes (p value 6.3E-4), while the B form increases the translation of genes in this category (p value 1.0E-7) (Fig. 3d). Analysis of the genetic and physical interactions of the top 35 translated genes in *ul30bb* cells identified a tight network of proteins involved in translation, large subunit (LSU) biogenesis, ribonucleoside synthesis, cell wall and transport (Fig. 3e). This network of interacting proteins is consistent with the B form-induced changes in translation and resistance to staurosporine, which is linked to cell wall integrity ^23, 24^. We conclude that *uL30* paralogs differentially modulate the translation of cell periphery genes, which may explain the paralog-dependent differences in staurosporine resistance.

### The minor paralog of uL30 tempers staurosporine-dependent modification of translation

Staurosporine altered the expression of ∼600-800 genes in wild type and *ul30aa* cells, while only affecting 150 genes in *ul30bb* (Fig. 3f-g). Furthermore, the only gene ontology category that was inhibited in wild-type and induced in *ul30bb* cells after staurosporine treatment was in cell periphery genes (Fig. 3h). Genes that are most upregulated by staurosporine in the presence of the B, and not the A form are also associated with the cell wall and membrane proteins (Fig. 3i). These data indicate that *uL30B* increases resistance to staurosporine by increasing the translation of cell periphery genes. We therefore hypothesized that the minor paralog of UL30 might affect the response to cell wall integrity drugs other than just staurosporine. As postulated, cells expressing the A form were sensitive, while those expressing the B version were resistant to the cell wall integrity drugs ketoconazole and caffeine (Extended Data Fig. 6). We conclude that uL30b-dependent resistance to drugs is not limited to staurosporine but extends to other drugs affecting the integrity of cell wall.

### *uL30* minor paralog selectively induces the translation of mRNA with long open reading frames

Comparing the features of the genes that are differentially translated by *uL30* paralogs identified clear differences in the length of the open reading frames. Coding regions of the top 20 genes, which are more translated in the presence of *uL30b,* are much longer than those that are under-translated (Fig. 4a). This difference in length was also observed in the presence of staurosporine (Fig. 4a). Comparing the length of differentially translated ORFs to the average ORF size also confirmed the preferential translation of long ORFs in cells expressing minor paralog (Fig. 4b). The ORF length-dependent translation was also validated by qRT-PCR using the model long ORF *GLT1* and short *EGD1* ORF (Fig. 4c). The differences in the translation of mRNAs of different lengths was not due to differences in the early initiation phases, as the amount of free mRNA did not vary in cells expressing different paralogs (Fig. 4d). Instead, we observe increased association of the long ORFs with the heavy polyribosome in cells expressing uL30B. Curiously, this minor isoform also decreased the co-sedimentation of the long ORF with the 60S fractions (Fig. 4d). It is not clear if this unusual co-sedimentation pattern represents abnormal association with the subunit or a coincidental co-sedimentation of an independent mRNA protein complex. In all cases, it is clear that uL30B increases translation of long ORFs, at least in part, through increased association with ribosomes.

**Figure 4.**
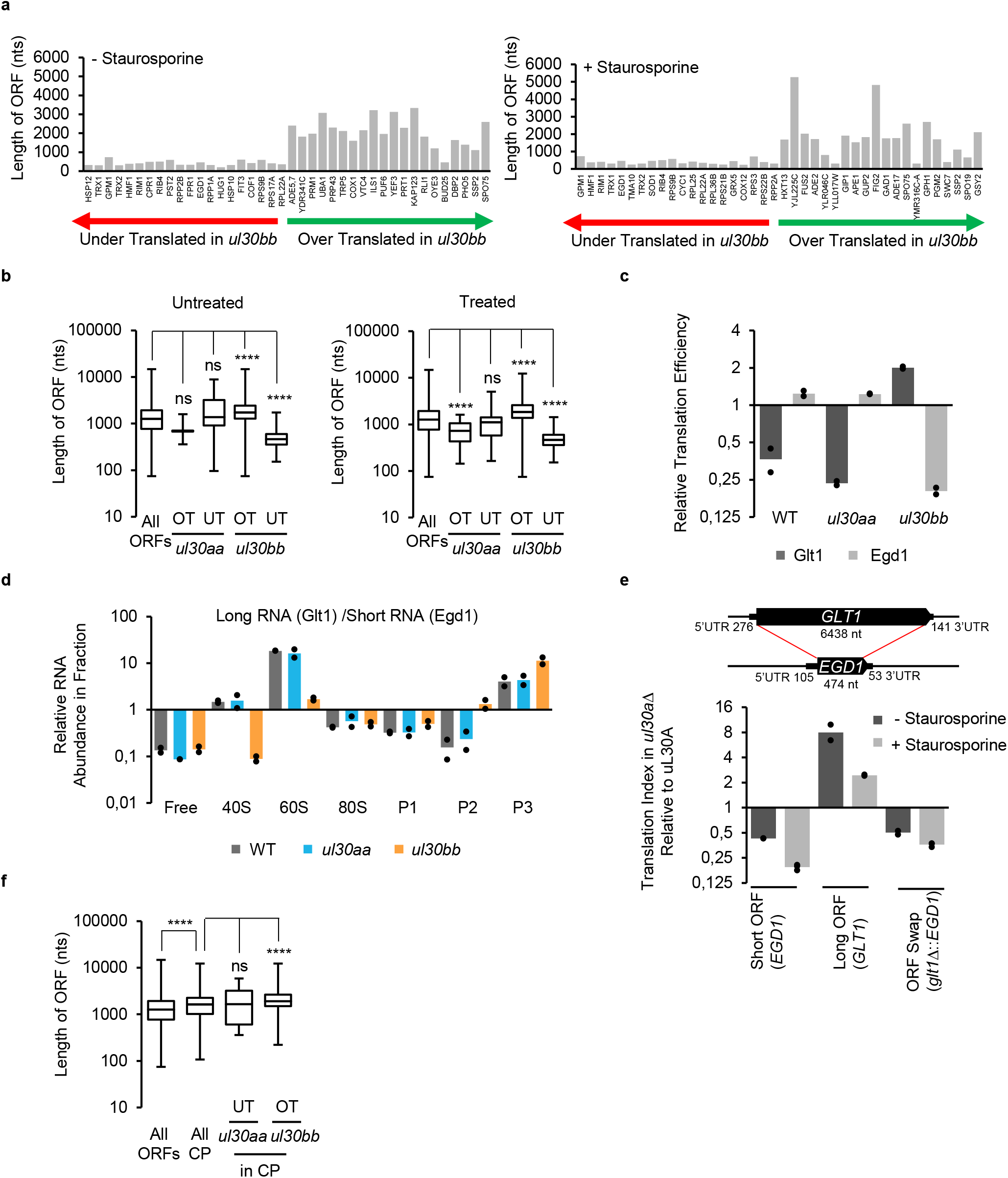
Paralogs of *uL30* modulate cell response to drug by altering the translation of genes with long open reading frames. **a,** The top 20 genes over- or under-translated in *uL30bb* strain in absence (left panel) or presence (right panel) of staurosporine are plotted relative to the length of their ORF in nucleotides. Genes are ordered on the X-axis as function of the magnitude of their change in translation. **b,** Box plots representing the length of ORFs for genes that are over- (OT) and under-translated-(UT) in *ul30aa* and *ul30bb* in absence (left panel) or presence (right panel) of staurosporine and compared to the ORF length of all yeast mRNA. Boxes limits represent the first and third quartiles, the middle line is the median and whiskers extend to data points limits (****p <0.0001 by Mann-Withney test). **c,** Translation efficiency of model mRNAs Glt1 (long) and Egd1 (short) examined using qRT-PCR for 2 biological repeats. **d,** The relative enrichment of long mRNA (Glt1) over short mRNA (Egd1) was monitored across sucrose gradients fractions using qRT-PCR for 2 biological repeats. (P1 are di-trisomes, P2 are quadra-penta-hexasomes and P3 are heptasomes and heavier polysomes fractions). **e,** The long ORF of *GLT1* was substituted with the short ORF of *EGD1* (top panel) in strains containing or lacking major paralog of *uL30* and translation of the chimeric mRNA examined in presence (+) or the absence (-) of staurosporine by RT-qPCR (lower panel). Graph shows the relative change in translation for the endogenous short gene (*EGD1*), long gene (*GLT1*) or chimeric gene (*glt1*Δ::*EGD1*) upon deletion of uL30A for 2-3 biological repeats. **f,** Length of ORFs coding for all cell periphery proteins (CP) and those that are over-(OT) or under-translated (UT) in *ul30aa* and *ul30bb* strains are represented in box plot as in (**b**).is compared to average length of all ORFs (****p <0.0001 by Mann-Withney test). RNA sequencing for panels **a,b** and **f** has been performed on 2 biological repeats.

To evaluate the ORFs length contribution to uL30B-dependent translation, we replaced *GLT1’s long* ORF with the short *EGD1* ORF and monitored translation before and after staurosporine treatment. Strikingly, changing the ORFs length abolished both paralog-and staurosporine-dependent translation confirming the importance of ORF length to each paralog’s effect on translation (Fig.4e). Interestingly, ORFs coding for cell periphery proteins, as a group, are longer than the average ORF length, explaining their increase expression in the presence of uL30B (Fig. 4f). We conclude that the *uL30* paralog-dependent response to staurosporine is mediated by differential translation of genes with long ORFs.

### The N-terminal domain of uL30 is required for resistance to staurosporine

To evaluate the contribution of the chromosomal loci and regulatory sequences to the paralog’s functional specificity, we compared cells expressing the cDNA of either paralog from identical plasmids and promoters (Fig. 5a-b). The cDNAs were transcribed from the promoter of the 40S subunit protein *eS28A/RPS28A* as the sole source of uL30 protein in the cell. In comparison to the native copy, plasmids equally increased the mRNA abundance of both paralog (Fig. 5c and Extended Data Fig. 7a). However as would be expected, the increased mRNA abundance did not increase the amount of uL30 proteins, as free RPs are rapidly degraded (Extended Data Fig. 7b)^25^. Importantly, plasmids did not alter the amount of other RPs either (Extended Data Fig. 7c-d). However, expression of the ORFs from the plasmid eliminated differences in the paralogs’ Kozak scores and thus reduced the preferential association of the A mRNA with heavy polyribosome (Extended Data Fig. 4). These results indicate that cells expressing *uL30* copies from plasmids generate equal amount of protein forms and ribosomes but differ from wild-type cells in paralog mRNA abundance translation pattern.

**Figure 5.**
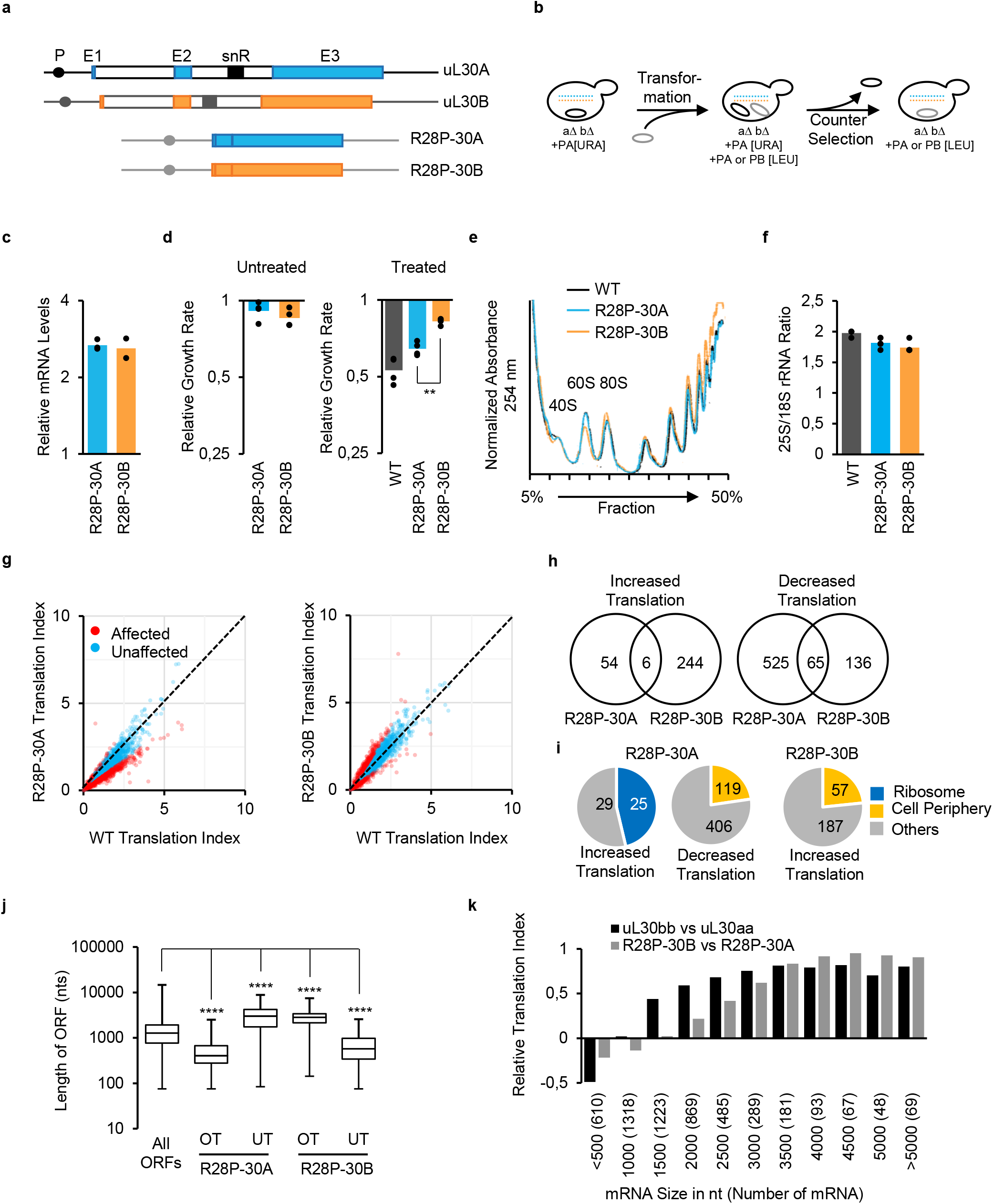
*uL30B* promotes translation of long mRNAs and staurosporine resistance independent of the non-coding regulatory sequences. **a,** Schematics of uL30 gene structure and constructs. Endogenous wild-type genes are shown on top and those of plasmids expressing paralogs cDNA from *RPS28A* promoter and *ADH1* terminator sequence are below. Blue, orange and white boxes indicate exons of *uL30A*, *uL30A* and introns, respectively. Lines show 5’ and 3’ UTRs. P, E1, E2, E3 and snR indicate position of promoter, exon 1-3, and snoRNA. **b,** Strategy for creating haploid yeast strains carrying deletion of both paralogs chromosomal copies and expressing either R28P-30A or R28P-30B plasmids using standard plasmid shuffling. **c,** uL30 mRNA expression from plasmid was quantified by RT-qPCR relative to endogenous uL30 for 2-3 biological repeats. **d,** Growth of cells harbouring R28P-30A or R28P-30B plasmids was compared to that of WT in absence (left panel) or presence (right panel) of staurosporine in 4 biological repeats (**p <0.01, by two-tailed paired t-test). **e,** Polysome profiles of cells expressing uL30 from plasmid. Curves shown are representative examples from 3 biological repeats. **f,** The ratio of 25S to 18S rRNA was determined by capillary electrophoresis for 4-5 biological repeats. **g,** Translation index of different mRNAs (expression >1 TPM) in cells expressing either paralog of uL30 from plasmid was compared to that of wild-type. mRNAs that were differentially translated (difference >log_2_ 0.5) in plasmid containing strains are shown in red. **h,** Venn diagrams of number of genes with changed association to polysomes identified in (**g**). **i,** Distribution of the number of genes present in enriched component gene ontology categories (p <0.001) for genes with specific changes in association to polysomes in cells expressing uL30 from plasmids. **j,** Box plots representing the length of ORFs for genes that are over- (OT) and under-translated-(UT) in cells expressing uL30 from plasmid. Boxes limits represent the first and third quartiles, the middle line is the median and whiskers extend to data points limits (****p <0.0001 by Mann-Withney test). **k,** Bar graph showing the relative translation index for the B/A paralogs according to mRNA sizes. RNA sequencing for panels **g-k** was performed on 2 biological repeats.

Comparison between cells expressing different versions of *uL30* genes from plasmids indicated, that unlike the chromosomal copies, the plasmid-borne versions equally supported growth and subunits production under normal conditions (Figure 5d-f). However, despite the lack of differences in ribosome biogenesis, cells expressing the B form from the plasmid were more resistant to staurosporine than those expressing the A form (Fig. 5d). The B form-specific increase in translation of long ORFs and cell periphery genes was also maintained in cells expressing the paralog from plasmids (Fig. 5g-j). Indeed, the translation index obtained with the chromosomal and plasmid copies were perfectly correlated (Fig. 5k). This indicates that the plasmid-dependent differences in RNA abundance and / or translation eliminate the differences in ribosome biogenesis without affecting the paralog specific resistance to staurosporine. We concluded that paralog-specific translation and drug resistance are driven by differences in protein sequence rather than differences in expression patterns.

### The paralog specific functions of *uL30* are driven by differential acetylation of its N-terminal domain

Most of the differences between the uL30 copies are found in the first 42 amino acids of the N-terminal domain (Fig. 6a). Four out of the five non-identical amino acids and 10/13 different sub-optimal codons are near the 5’end. Furthermore, the paralog’s N-terminal residues are differentially acetylated. The alanine residue of the B form is 100% acetylated, while the serine residue at the corresponding position of the A version is mostly unmodified (Fig. 6a-b). All proteins produced in cells lacking the *uL30A* gene (*ul30bb*) were 100% acetylated, while those lacking the *uL30b* paralog (*ul30aa*) only produced unmodified proteins (Fig. 6b). The acetylated protein was more abundant in heavy polyribosomes, and exposure to staurosporine increased in the amount of ribosome incorporating the 100 % acetylated B form (Fig. 6c and Fig. 2g). Paralog acetylation depends on the N-terminus sequence, since swapping the first 42 amino acids of the B form with the A version abolished the hyper-acetylation of the B form (Fig. 6d). Remarkably, swapping the N-terminal domains on plasmids or the chromosomal copies reversed the effects of the paralog on translation and on resistance to staurosporine (Fig. 6e and Extended Data Fig. 8). The resistance to staurosporine was also reversed when only the amino acids differentiating the A and B forms were swapped (Fig. 6f). We conclude that staurosporine resistance is driven by acetylation of specific amino acid residues in the minor uL30 paralog.

**Figure 6.**
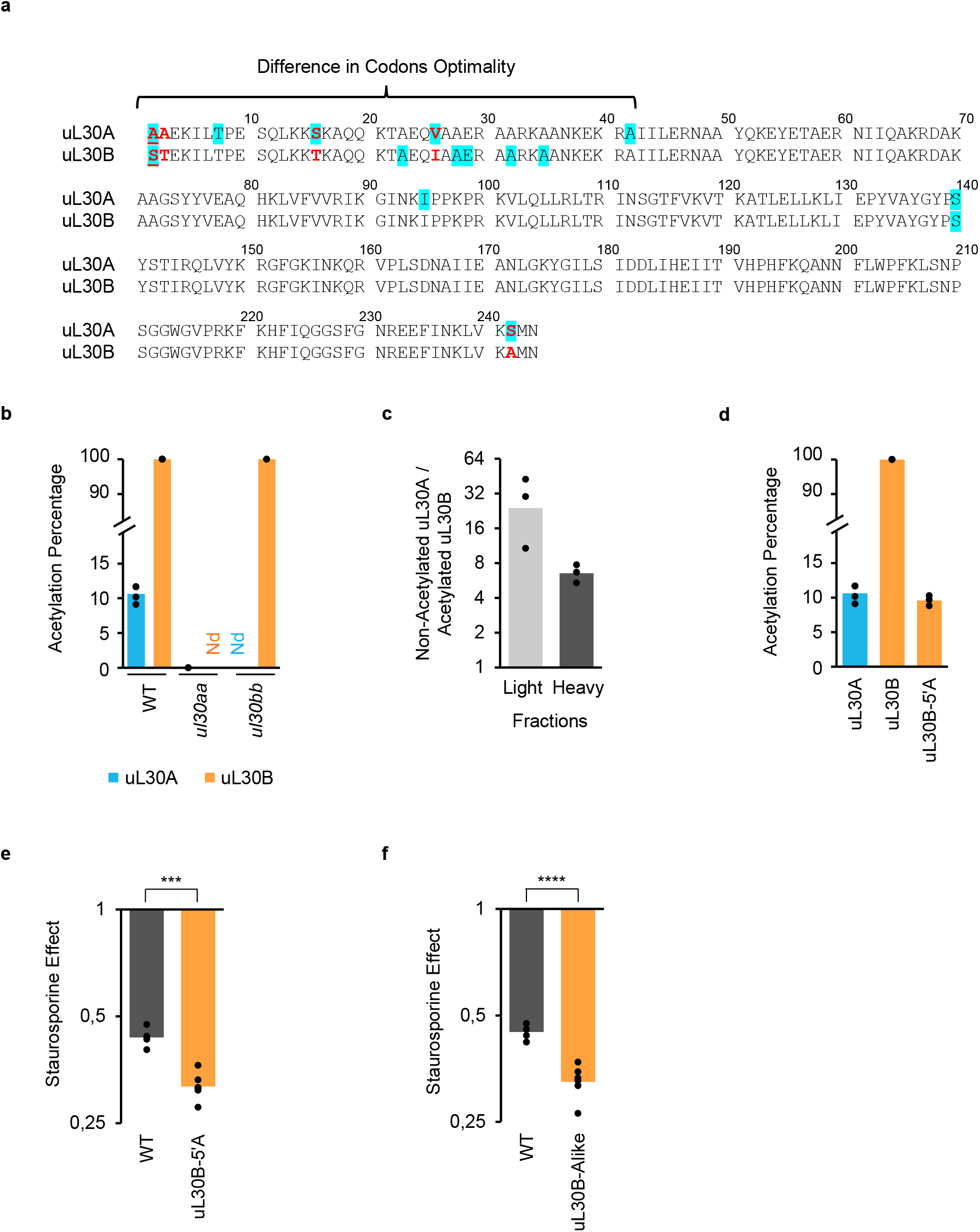
Post-translation modification alters function of uL30 paralogs. **a,** Amino acid sequences of uL30 paralogs were aligned using Clustal/W2. Different amino acids are indicated in red, amino acids encoded by suboptimal codon are highlighted in cyan and position of N-terminal acetylation is underlined. **b,** N-terminal acetylation was measured by mass-spectrometry in wild-type, *ul30aa* and *ul30bb* cells in 2-4 biological repeats and percentage of detected acetylated N-terminal peptides is reported. **c,** N-terminal acetylation level of uL30 proteins detected in light and heavy polyribosome fractions in WT cells using mass-spectrometry in 3 biological repeats. The ratio of non-acetylated uL30A over acetylated uL30B in each fraction is reported. **d,** N-terminal acetylation was measured by mass-spectrometry in wild-type and uL30B-5’A mutant for 3-4 biological repeats and percentage of detected acetylated N-terminal peptides is reported. **e,f,** Relative growth rates of WT and uL30B-5’A mutant (**e**) or ul30B-Alike mutant (**f**) after staurosporine treatment for 4-7 biological repeats. (***p<0.001, ****p<0.0001 by two-tailed t-test assuming unequal variance).

## Discussion

In this study, we show that exposure to drugs modifies ribosomes composition and translation by altering the ratio of proteins generated from duplicated ribosomal protein genes (dRPG). Most yeast ribosomes are generated from one predominant (major) housekeeping paralog required for normal growth and a minor paralog needed under certain conditions (Fig. 1). Interestingly, we discovered that quasi-identical RPGs could produce two differentially modified proteins with different effects on translation. For the uL30 gene-pair the major paralog is hypo-acetylated and required for optimal ribosome biogenesis and cell growth, while the minor form is hyper-acetylated to provide optimal drug resistance (Fig. 2, 4 and 6). Paralog-dependent differences in drug-resistance originates from variation in the sequence and post-translation modification of the protein’s N-terminal domain (Fig. 6). Together our data provide clear evidence of programmed growth condition-specific changes in ribosome composition that promote drug resistance by modifying translation.

The paralog-specific effects of *uL30* include differences in the 60S biogenesis, translation and staurosporine resistance. The paralog specific contribution to the synthesis of the 60S subunit is linked to the paralogs’ chromosomal loci, while the differences in translation and drug-resistance are linked to differences in the modification and sequence of N-terminal amino acids. The different effects of the paralogs on ribosome biogenesis are consistent with the role of *uL30* in assembling the 25S rRNA domains ^26^. Furthermore, the paralog-specific effects appear to be independent of the gene copy number (Extended Data Fig. 2). The experiments in Fig. 5 as well as previously reported intron deletions indicated that the difference in uL30 paralog functions is not due to their introns, or the snoRNAs embedded therein ^27^. Instead, uL30B protein appears to be less efficient than uL30A in supporting ribosome biogenesis, a defect that could be overcome by increasing the expression of uL30B from plasmids (Fig. 2c and 5e). Expression of uL30B from a plasmid may temper the defect observed with the chromosomal copy in ribosome biogenesis by modifying the timing and location of the gene expression. Natural *uL30* paralogs have different localization patterns ^28^ and overexpression of these paralogs may eliminate this difference through mislocalization, which in turn may blur their effects on ribosome biogenesis.

In contrast to ribosome biogenesis the second category of paralog-specific effect reported here, namely differences in translation and resistance to drugs affecting cell wall integrity, mostly stems from differences in the paralog sequences and consequent modification of their N-terminal domains. The N-terminal domains of the two paralogs, which differ by 4 amino acids, are predicted to form different structures and they are differentially acetylated (Fig. 6a and Extended Data Fig. 2b). Furthermore, exchanging the paralog N-terminal amino acid sequence or mutation of the differentially acetylated amino acid abolishes the paralog-specific effects on translation and drug resistance (Fig. 6d-f and Extended Data Fig. 8). Some differences in posttranslational modification stem from differences in the paralog translation rate (Extended Data Fig. 4), which may alter their amenability for posttranslational modification. This type of differential modification has the potential to substantially increase the ribosome diversity, as most ribosomal proteins are subject to a variety of posttranslational modifications ^12, 29, 30^.

Together the data presented in this study point to a model of dRPG function in which the major paralog is the most efficiently expressed gene under normal condition enabling it to carry the housekeeping functions, while the minor paralog serves an auxiliary role that become important in responding to stress (Fig. 7). In the case of uL30, under normal conditions, the gene coding for the A form is more efficiently translated and less posttranslationally modified than the B from. The increased production of the A form ensures optimal ribosome biogenesis resulting in the accumulation of ribosomes containing the A form.

**Figure 7.**
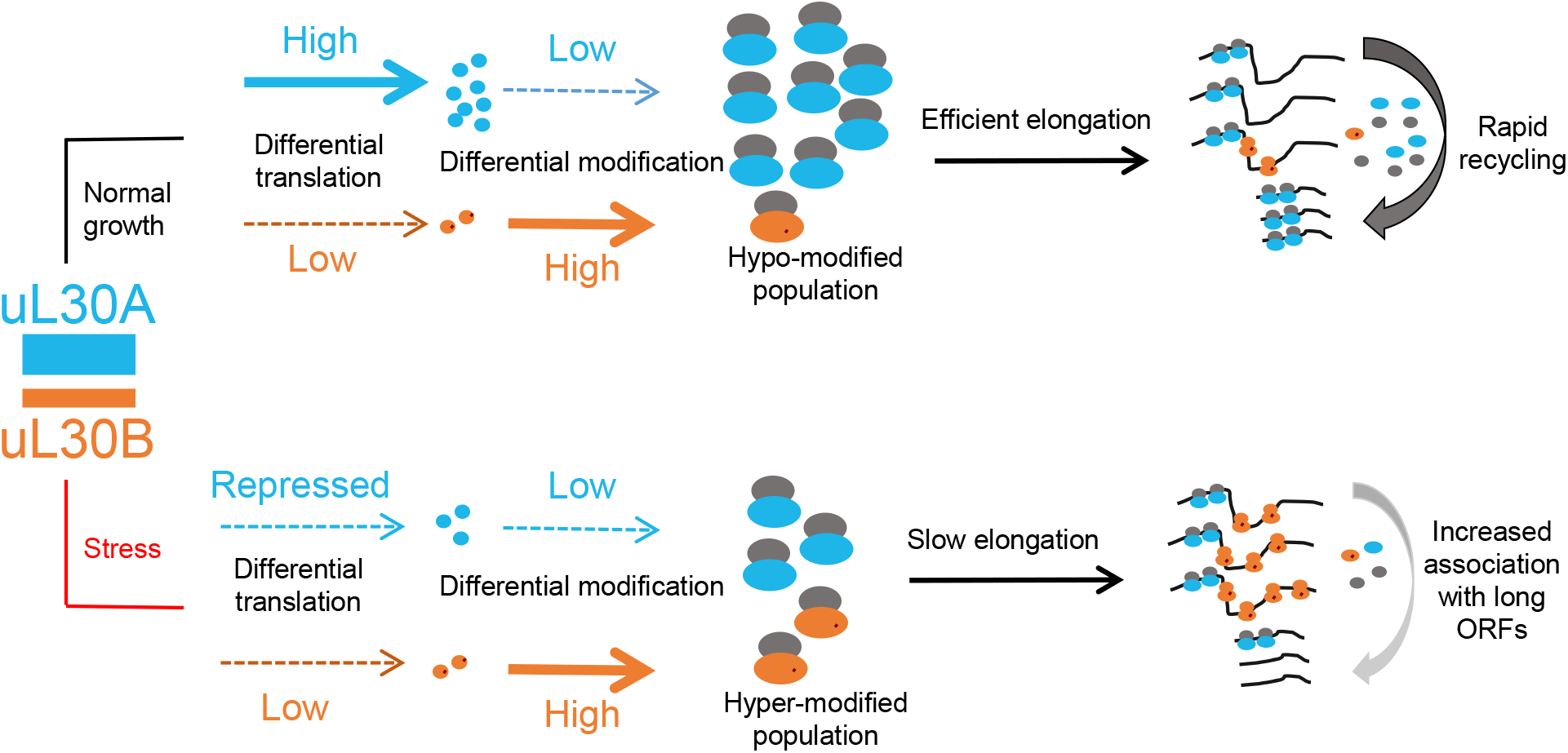
Schematic representation of paralog dependent translation modulation. Expression of major and minor paralog and their impact on translation is depicted under normal growth condition or after exposure to stress (e.g., staurosporine). Major (uL30A) and minor (uL30B) paralogs are indicated as cyan and orange boxes, respectively. Large ribosomal subunits incorporating major and minor paralogs are indicated as blue and orange ovals. Post-translational modification of the minor paralog is indicated as red dots and mRNAs are indicated as black lines.

In this model, the abundance of hypo-acetylated uL30-A form ribosome ensures optimal translation, especially of short mRNAs, which in turn ensures the most rapid elongation and recycling rate. In contrast, translation is inhibited under stress, reducing translation of the major paralog, with little effect on the already under-translated minor version. As a result, more ribosomes containing the hyper-acetylated minor protein form, slowing-down translation further and increasing the association of ribosomes with long mRNAs. In this model then, the paralog changes the translation equilibrium of long and short ORFs. This switch is likely due to reduced availability of free 60S subunits. Either through reduced production of the 60S or delayed release from the mRNA. This model is corroborated by the experimental cells expressing the minor paralog either from the chromosomal copy where there reduced amount of 60S or from plasmids where mRNAs are more associated with polyribosomes. In all cases, the effect is clearly linked, to the kinetics, and or pattern, of expression and modification of ul30 paralogs as the expression from plasmids alters the effects observed with chromosomal copies. Together this work shows that the observed effect is not due to differences in the capacity of the paralogs to generate adequate dose of ribosomal protein but depends on a combination of paralog-specific expression pattern, amino acids sequence and modifications.

## Methods

### Experimental Model

*Saccharomyces cerevisiae* strains used in this study share the BY background ^31^ and are described in Supplementary Table 1. Cultures were grown in yeast complete (YC) media (1.7 g/l yeast nitrogen base without amino acids and ammonium sulfate, 1 g/l sodium glutamate, 100 mg/l cysteine and threonine, 85 mg/l tryptophan, 80 mg/l leucine, 60 mg/l lysine, 50 mg/l aspartic acid, isoleucine, methionine, phenylalanine, proline, serine, tyrosine, uracil and valine, 20 mg/l adenine, arginine and histidine, pH 6) with 2 g/l dextrose at 30°C. Strains were constructed using standard procedures ^32^ and gene replacements verified by PCR using oligonucleotides described in Supplementary Table 2. Plasmids R28P-30A and R28P-30B were synthesized and sequenced by Biobasic.

### Growth Assays

Overnight saturated starter cultures from freshly streaked single colonies were used to inoculate precultures grown to reach a cell density of 1.67-3.33E7 cells/ml after an overnight incubation. These precultures were diluted the next morning to 0.83-1.25E6 cells/ml and the experiment started when the cultures reached 2.33-2.67E6 cells/ml. For growth assays in tubes, IC 50 (drug concentration which reduces the growth of wild-type cells by 50%) quantities of drugs were added at this point and absorbance at 660 nm was measured at regular intervals for 10-48 hours. For growth assays in 96-well microplates, the absorbance at 660 nm of 100 µl cultures was measured at 10 minutes intervals for 48 hours in a Biotek Powerwave or RNAEpoch2 readers. The maximal growth rates (doubling times) for each strain were calculated as previously described ^33^.

### Total RNA Isolation

Cells from 5-10 ml cultures grown to 1-1.33E7 cells/ml were collected by centrifugation, washed with DEPC-treated water, and resuspended in 300 μl of LETS buffer (0.01 M Tris-HCl pH 7.5, 0.1 M LiCl, 0.01 M Na_2_EDTA pH 8.0, 0.2% SDS), then 300 μl of phenol and 1 volume of acid-washed glass beads were added. Cells were broken by 10 cycles of vortexing for 30 seconds and cooling on ice for 30 seconds. Proteins were removed by phenol:chloroform extraction and the RNA precipitated by the addition of 2.5 volumes of ethanol and 0.3 M sodium acetate pH 5.5. RNA samples were resuspended in 50 μl of DEPC-treated H_2_O and quantified by Nanodrop. Samples diluted to 25-50 ng/µl were used for 25S/18S quantification on an Agilent TapeStation 2200 using RNA screen tape.

### Polysome and RNA Preparation for Translation Index Analysis

Polysomes were prepared as described by ^34, 35^ with the following modifications. 50 ml pre-cultures made by inoculating with a single colony from a freshly streaked plate and incubated overnight to reach a cell density of 5.83-8.33E7 cells/ml. The precultures were diluted to 0.83E6 cells/ml in a volume of 400 ml. Staurosporine treatment started when the cultures reached a density of 3.33E6 cells/ml and ended when the cultures reached a density of 1-1.33E7 cells/ml. The cells were collected in precooled 400 ml plastic bottles, cycloheximide was added to a final concentration of 0.1mg/ml and they were immediately spun down at 3000 g for 5 min at 4°C. The pellets were washed twice in cell lysis buffer (20 mM Tris-HCl pH 8, 140 mM KCl, 1.5 mM MgCl_2,_ 0.5 mM DTT, 0.1 mg/ml cycloheximide, 1% Triton X-100, 1 mg/ml heparin). The washed cells were resuspended in 3 ml of lysis buffer, 500 µl of acid washed glass beads were added, and the samples were vortexed 4 × 20 seconds at full speed on a vortex Genie 2 (VWR Scientific Products). The cellular debris were removed by centrifugation at 2,600g for 5 min at 4°C and 1.5 ml of the lysate was transferred to a micro-tube and centrifuged again at 9500 g for 5 min at 4°C. Quickly (within 2.5 hours) the supernatant was layered onto a 5-50% sucrose gradient prepared with 20 mM Tris-HCl pH 8, 140 mM KCl, 5 mM MgCl_2_, 0.5 mM DTT, 0.1 mg/ml cycloheximide, 1 mg/ml heparin on Gradient Master 108 (Biocomp) at an angle of 60° for 5 minutes at 25 rpm and placed at 4°C for at least 10 hours) and centrifuged at 34,000 g for 13 hours in a Beckman SW28 rotor at 4°C. Gradients were fractionated using a Teledyne Isco gradient collector with pump speed set at 2 ml/min and chart speed at 30 cm/h. The fractions were collected in 13 ml round bottom polypropylene tubes and transferred to 50 ml Nalgene Oakridge centrifuge tubes (3118-0050). Two volumes of guanidine-HCl 8 M and 3 volumes of 100% ethanol were added to samples for an overnight crude RNA precipitation at −80°C. The samples were centrifuged at 20,000 g for 20 min at 4°C. The pellets were washed with 3 ml of 85% ethanol and spun again for 20 min at 20,000 g at 4°C. The pellets were dissolved in 3 ml of DEPC-treated water by vigorously vortexing 4 × 20 seconds. The resuspended RNAs were transferred to 13ml polypropylene round-bottom tubes and precipitated with 300 µl of 3 M sodium acetate pH 5.3 and 2.5 volumes of 100% ethanol overnight at −80°C and then centrifuged at 20,000 g for 20 min at 4°C. The pellets were washed with 3 ml of 85% ethanol and spun again for 20 min at 20,000 g at 4°C. The pellets were then dissolved in 1 ml of DEPC-treated water by vigorously vortexing 4 × 20 seconds. Proteins were removed by successive phenol:chloroform and chloroform extractions. 900 µl of aqueous phase was transferred to a 2 ml microtube and RNAs were precipitated with 1 ml of 3 M LiCl overnight at −80°C. The RNAs were precipitated in a microcentrifuge at 13,000 g for 20 min at 4°C and pellets were washed with 85% ethanol and spun again for 20 minutes at 13,000 g at 4°C. The pellets were resuspended in 350 µl of DEPC-treated water. To remove LiCl, samples were precipitated again with 35 µl 3M sodium acetate pH 5.3 and 1,155 ml of 100% ethanol overnight at −80°C. The pellets were washed with 75% ethanol and dried in a speedvac at 40°C for 20 minutes. The pelleted and dried pure RNAs were finally resuspended in 100 µl of DEPC-treated water and stored at −20°C before deep sequencing analysis and RT-qPCR.

### RNA Sequencing

Starting with 1 µg RNA from total, light polysome (40S+60S+80S) and heavy polysome (tetrasome and heavier) fractions, we enriched mRNA using the NEBNext Poly(A) mRNA Magnetic Isolation Module (E7490S). NGS libraries were then prepared with NEBNext using Ultra II Directional RNA Library Prep Kit for library construction in a poly(A) mRNA enrichment workflow (E7760S) according to the manufacturer’s protocol. Individual libraries were purified with Ampure XP beads, analysed on an Agilent Bioanalyser 2100 (HS DNA) for size and quality then quantified using Qubit 2.0. A first round (wild-type and *uL30bb* for light and heavy polysome fractions, untreated and staurosporine treated samples) of sequencing was performed on Illumina’s NextSeq sequencer. Two pools (one for each N) of 8 libraries/pool were sequenced in 2 separate runs using NextSeq 500/550 High output v2 kit (150 cycles) FC-404-2002 achieving a mean depth of 70 million reads per sample. A subsequent round (Wild-type and *uL30bb* total extracts; *uL30aa* total light, and heavy polysome fractions) of sequencing was performed with a pool of 20 samples for a mean depth of 30 million reads per sample. Two biological replicates of WT, R28P-30A and R28P-30B were run similarly as a pool of 18 samples.

### RT-qPCR and Droplet Digital PCR (ddPCR)

Contaminant genomic DNA from 6-25 μg of RNA, extracted either from total extracts or from sucrose gradient fractions, was removed by treatment with 33 units of Qiagen RNAse-free DNase (79254) on Qiagen RNeasy Minikit (74106) spin columns for 25 min at 37°C. DNase was inactivated and washed away by the provided buffer and RNAs were eluted twice with 70°C DEPC-treated water. A total of 50 ng DNase-treated RNA was used as template for reverse transcription using either RT Transcriptor from Roche and random hexamers or Moloney Murine Leukemia Virus-RT (MMULV-RT) locally produced by the protein purification service of Université de Sherbrooke. The qPCR reactions were carried out in 384 well plates in a Biorad CFX384^TM^ Real Time System thermocycler in 10 μl volumes using the BioRad iTaq Universal SYBR Green Supermix, 1 ng of cDNA and 200 nM of primers described in Supplementary Table 2. In Fig. 4d, 2 pg of *E.coli* rRNA (Roche) was added as spike-in to equal volumes of RNA extracted from sucrose gradient fractions for the reverse transcription. Droplet digital PCR reactions were carried in duplex using QX200 ddPCR Supermix (Bio-Rad) and 0.06 ng of cDNA in presence of 250 nM of the labeled probes and 900 nM of the primers ^36^. The mixes were processed in a Bio-Rad QX-200 droplet generator. PCR reactions were carried in a C1000 deep well Thermocycler (Bio-Rad), then the plates were transferred to a QX200 reader (Bio-Rad). The concentration in copies/ul was obtained using the QuantaSoft (Bio-Rad).

### Protein Preparation for Mass-Spectrometry

Protein quantification by mass spectrometry was carried as previously described by ^37^ with the following modifications. Cells were grown and harvested as described in the polysome preparation section. For Fig.1c, 2f and Extended Data Fig. 2c,g, 5e and 7b,d, the proteins were extracted in protein extraction reagent type 4 in 50 mM tris pH 8.0. For Fig. 2g and Extended Data Fig. 2d cells were grown and harvested and the sucrose gradients were performed as described in the polysome preparation section. To avoid interference, no heparin was added in extraction buffer nor in sucrose gradients. Spectra (3500 Da, Spectrum laboratories) tubing was used to dialyse the protein extracts against TSM buffer (10 mM Tris, 3 mM succinic acid, 10 mM MgCl_2_, pH 8) over 70 hours at 4°C with continuous agitation and 6-8 changes of 5 litres of buffer. The dialysed samples were frozen at −80°C in 50 ml Nalgene Oakridge centrifuge tubes (3118-0050), lyophilized at low atmospheric temperature (150-250 mtorr) and resuspended in phosphate buffer (140 mM NaCl, 2.5 mM KCl, 4 mM Na_2_HPO_4_, 1.5 mM KH_2_PO_4_, pH 7.3). The samples were mixed by slow agitation at 4°C with 0.4 volume of 1M MgCl_2_ and 2 volumes of glacial acetic acid, then centrifuged at 20,000 g for 10 minutes at 4°C. A second dialysis was performed against 0.5 % acetic acid over 70 hours at 4°C with continuous agitation and 6-8 changes of 5 litres of buffer and lyophilized. Paralog specific protein quantification was performed on LC-MS/MS TripleTOF 5600 mass spectrometer (ABSciex; Foster City, CA) equipped with a DuoSpray source at PhenoSwitch Bioscience (Sherbrooke, Canada). Reagent 4 (6 M urea, 2 M thiourea, 4% CHAPS in 50 mM Tris pH 8) was used to suspend lyophilized proteins and Pierce 660 protein assay used to estimate the protein quantity. DTT was used to alkylate and reduce 40 µg of protein followed by overnight precipitation with 1 volume of ice-cold methanol and 8 volumes of ice-cold acetone at −80°C. After centrifugation, pelleted proteins were washed 3 times with ice-cold methanol and briefly air dried. A first digestion was performed using 1:30 w/w ratio, of lysine C to proteins, followed by digestion using the same ratio of trypsin in 0.75 M urea and 50 mM Tris pH 8 for 4 hours at 37°C. Formic acid 2% v/v was used to stop digestion followed by peptide purification by solid-phase extraction on a polymeric reverse-phase cartridge (Phenomenex, Torrance, CA). LC-MS/MS. Digested and purified proteins were ionized by an ESI (Electron spray ionization) source in a 25 µm ESI probe (Eksigent) using a microLC200 system (Eksigent) equipped with a 150 mm x 300 µm HALO C18 2.7 µm column (Eksigent). Samples were injected by overfilling a 5 µl injection loop. A gradient of water containing 0.2% formic acid and 3% DMSO and ethanol containing 0.2% formic acid 3% DMSO were used to perform chromatography at 50°C. Rolling collision energy and optimized SWATH windows in positive product ion mode with a mass range from 100 to 1800 m/z and in high sensitivity MS/MS mode were used for data acquisition. For Fig. 6b-d, proteins were extracted or resuspended in 8 M urea, 20 mM Tris pH 8. Samples containing 50 µg total proteins were treated with 5 mM DTT for 2 minutes at 95°C, cooled down to RT for 30 min then treated with 7.5 mM chloroacetamide at RT for 20 min in the dark. Following dilution with 3 volumes of 50 mM NH_4_HCO_3_, samples were digested overnight at 37°C with 1 µg of LysC (Promega) ^38^. Samples were purified and desalted on 100 µl ZipTips (Pierce), eluted in 1% formic acid / 50% acetonitrile, dried and resuspended in 30 µl of 1% formic acid. Following quantification at 205 nm, 1.5 µg of peptides were separated at the Plateforme de Protéomique (Université de Sherbrooke, Canada) on a Dionex Ultimate 3000 nanoHPLC system using an Acclaim PepMap100 C18 column (0.3 mm id x 5 mm, Dionex) followed by an EasySpray PepMap C18 nano column (75 µm x 50 cm, Dionex) with a 5%-35% of 90% acetonitrile / 0.1% formic acid over 240 min at a rate of 200 nl/min. Samples were ionized by an EasySpray source and transferred to an OrbiTrap Q Exactive mass spectrometer (Thermo Fisher Scientific). The spray voltage was set to 2.0 kV and the column temperature to 40°C. Full scan MS survey spectra (*m/z* 350-1600) in profile mode were acquired in the Orbitrap with a resolution of 70,000 after accumulation of 1,000,000 ions. The ten most intense peptide ions from the preview scan in the Orbitrap were fragmented by collision-induced dissociation (normalized collision energy 25% and resolution 17,500) after the accumulation of 50,000 ions. Maximal filling times were 250 ms for the full scans and 60 ms for the MS/MS scans. Precursor ion charge state screening was enabled and all unassigned charge states as well as singly, 7 and 8 charged species were rejected. The dynamic exclusion list was restricted to a maximum of 500 entries with a maximum retention period of 40 s and a relative mass window of 10 ppm. The lock mass option was enabled for survey scans to improve mass accuracy. Data files were generated by the Xcalibur (v4.3.73.11) software.

### Total Protein Extraction and Western Blot Analysis

Cells from exponentially growing 40 ml cultures were pelleted by centrifugation for 3 min at 3000 g at 4°C and washed with cold water. The pellets were resuspended in 500 ul of lysis buffer (20 mM Tris pH8, 150 mM NaCl, 0.1% Triton X-100, 1 mM PMSF, 1 mM benzamidine, 1 μg/ml aprotinin, 1 μg/ml leupeptin, 1 μg/ml pepstatin A, 1 μg/ml antipain) and transferred to 2 ml tubes with 300 µl acid-washed glass beads. Cells were broken by 5 cycles of vortexing in a Bertin Precellys 24 homogenizer at 5000 rpm for 30 seconds followed by 30 seconds on ice. Lysates were cleared by centrifugation at 13,000 g for 15 minutes at 4°C and dosed by Bradford. Protein samples (20 µg) were separated on 15% SDS-PAGE and transferred to Protran nitrocellulose membranes (GE Healthcare). Membranes were blocked overnight at 4°C with 5% milk in TBS-T (20 mM Tris pH 7.6, 150 mM NaCl, 0.1% Tween-20), then incubated overnight at 4°C with the following primary antibodies: rabbit anti-L7 (Bethyl Laboratories, cat#A3100-741A-M, 1:1000), rabbit anti-L5 (kindly provided by Woolford lab, 1:2000), mouse anti-L3 (kindly provided by the Warner lab, 1:5000) and mouse anti-Pgk1 (Invitrogen/Thermo Fisher, cat#459250, 1:10,000) diluted in blocking solution. After washing with 4 changes of TBS-T over 40 minutes, the membranes were incubated for 90 minutes at room temperature with secondary antibodies (donkey anti-rabbit HRP-IgG, GE Healthacare, cat#NA934V, 1:5000 or goat anti-mouse HRP-IgG, GE Healthcare, cat#NA931V, 1:2000) diluted in blocking solution and washed as before. Western Lightning Plus-ECL reagents (Perkin Elmer) were used for bioluminescence and a LAS4000 system (GE Healthcare) for data acquisition. The Quantity One software from Bio-Rad was used for the quantification.

### Quantification and statistical analysis

#### RNA Sequencing Analysis

Saturation and coverage tests were performed for all samples and coverage was found adequate in all cases. Sequencing reads were aligned on a genome reference sequence (R64/sacCer3) using the STAR aligner (default parameters) and all valid positions mapped were kept. Gene expression was quantified in TPM for the sequence generated from total RNA extracts or CPM for sequence used for the calculation of the translation index using featuresCount from the Rsubread package ^39, 40^. A summary of the sequencing data is presented in Supplementary Table 3.

#### Mass Spectrometry Analysis

Data integration and analysis were initially performed by Peakview version 2.2. Peptide quantification carried out by Peakview was used for inter-sample normalization. Peak selection, protein quantification was performed using Skyline (MacCoss Lab). Relative paralog specific quantification was performed without any normalization (Fig. 1c, 2f-g and Extended Data Fig. 2c,d,g), expression of uL30 from plasmids was normalized to large subunit RPs (Extended Data Fig. 7b), while protein quantification of genes over- and under-translated in *ul30bb* and RP quantification was done after normalization to Pgk1 protein (Extended Data Fig.5e and 7d). For Fig. 6b-d, the raw files were analyzed using MaxQuant (v2.0.1.0) and Uniprot *S. cerevisiae* database (31/05/2021, 6079 entries). The settings used were: enzyme: LysC (K not before P); miscleavages: maximum of 4; fixed modifications: carbamidomethylation on C; variable modifications: carbamylation on K and N-termini, methionine oxidation and N-terminal acetylation; precursor ion mass tolerance of 7 ppm; fragment ion mass tolerance of 20 ppm; PSM FDR, protein FDR and site decoy fraction were set at 0.05. Peptides with the “Reverse” or “Potential contaminant” attributes were removed from the analysis. The intensities of the N-terminal peptides specific for uL30A and uL30B were summed and classified as N-terminal acetylated or not. Percentage of acetylation or paralog ratios were calculated within samples, hence no normalization was applied.

#### RT-qPCR Analysis

RT-qPCR were done in 2-3 biological replicates with 3 technical replicates. Relative quantification of mRNA was calculated according to the formula: RQ = 2^(CT_Reference_-CT_Test_)_Gene_/2^(CT_Reference_-CT_Test_)_Control_, where CTs are the cycle threshold values determined by the Bio-Rad CFX Manager software; Reference and Test represent strain, conditions or fractions compared; and Gene and Control represent the gene of interest quantified and the normalization control gene. Inter-sample comparison was achieved by normalizing mRNA to Osh6 mRNA or Nme1 noncoding RNA as internal control except for Fig. 4d where *E.coli* 23S rRNA spike-in was used to normalize across fractions. The translation index was calculated by dividing the amount of mRNA in heavy polysome fractions (tetrasome and heavier fractions) by the amount of mRNA detected in light polysome fraction (40S, 60S, 80S).

#### Gene Ontology Analysis

The Gene Ontology (GO) Component analyses were carried out using the SGD server (https://www.yeastgenome.org/) with the default settings of GO Term Finder. Only the major categories of component gene ontology with significant p-value < 0.001 using Bonferroni correction were considered.

#### Kozak Score Analysis

The Kozak score for 6717 transcripts were obtained by creating a matrix of the nucleotide frequencies (incidence of a nucleotide) for the six nucleotides upstream and three nucleotides downstream of the initiation codon AUG. A maximum score of 3.992 for the sequence ACAAAA-AUG-GCU was calculated.

#### Statistical Analysis

Statistical tests and Ns are specified in figure legends and were performed using GraphPad Prism for Windows. Significance is given as p-value from a t-test for growth rates or mRNA and protein level comparisons, or from a Mann Whitney test for genome-wide distribution comparisons as described in the figure legends.

### Data availability

RNA-seq data generated in this study have been submitted to the NCBI Gene Expression Omnibus (GEO; https://www. ncbi.nlm.nih.gov/geo) under the accession number GSE133457. Proteomic data presented in this study has been submitted to the Peptide Atlas (http://www.peptideatlas.org/) under accession number PASS01404. All strains are available upon request. Source data are provided with this paper. All other data supporting the findings of this study are available from the corresponding author on reasonable request.

## Acknowledgments

We thank Jules Gagnon for help with the computational analysis of mRNA feature, Camille Martenon-Brodeur and and Kloé Neszvecsko for help with the preparation of yeast strains and the initial screen for paralog specific effects, Julie Parenteau for advice and help with yeast genetics, Sonia Couture for help with RNA sequencing, Vincent Boivin and Gaspard Reulet for the analysis of sequencing data. Sequencing was performed in the U de S Rnomics Platform. Proteomic analysis was performed at Phenoswitch and U de S Proteomic Platform. We thank John Woolford for providing anti-L5 antibodies and Jonathan Warner for providing anti-L3 antibodies. This work was supported by NSERC and a Research Chair in RNA Biology and Cancer Genomics (S.A.E.).

## Author information

### Affiliations

RNA Group, Département de Microbiologie et d’Infectiologie, Faculté de Médecine et des Sciences de la Santé, Université de Sherbrooke, Sherbrooke, Quebec, Canada

Mustafa Malik-Ghulam, Mathieu Catala, & Sherif Abou Elela

RNA Group, Département de Biochimie et Génomique Fonctionnelle, Faculté de Médecine et des Sciences de la Santé, Université de Sherbrooke, Sherbrooke, Quebec, Canada

Michelle Scott

### Contributions

M.M.G. and M.C. designed and performed experiments, analyzed data, produced figures and participated in the writing of the paper. M.S. supervised the analysis of the RNA sequencing data and participated in the writing of the paper. S.A. proposed and designed experiments, analyzed data and wrote the paper.

### Corresponding author

Correspondence to Sherif Abou Elela.

---

**Extended Data Fig. 1.**
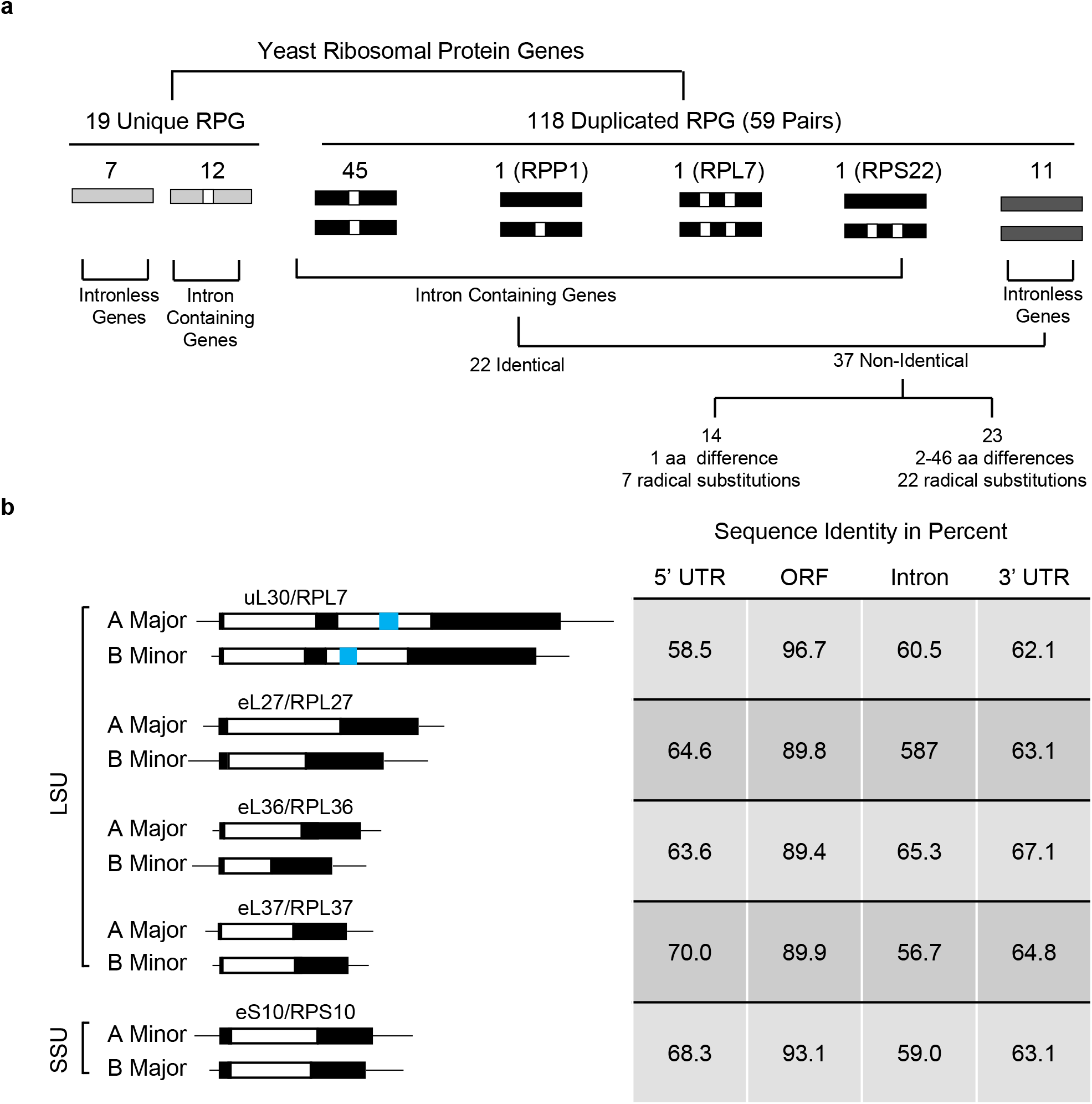
Classification and characteristics of yeast ribosomal protein genes. **a,** Schematic representation of yeast RPGs. RPGs were separated based on gene duplication, number of introns and number of amino acid difference between ohnologs. Non-identical paralogs contain at least one amino acid difference. Radical substitution indicates change in the amino acid that may cause charge or structural change. Black boxes indicate exons and white boxes indicate introns. **b,** Characteristics of the RPGs used to generate homogenized yeast strains. Genes coding for large (LSU) and small ribosomal subunits (SSU) proteins are indicated on the left. Both classical and universal protein names are indicated on the top of each gene pairs. Black, blue and white boxes indicates, exon, snoRNA and introns, respectively. The percent sequence identity between the different regions of each pair of genes is indicated on the right.

**Extended Data Fig. 2.**
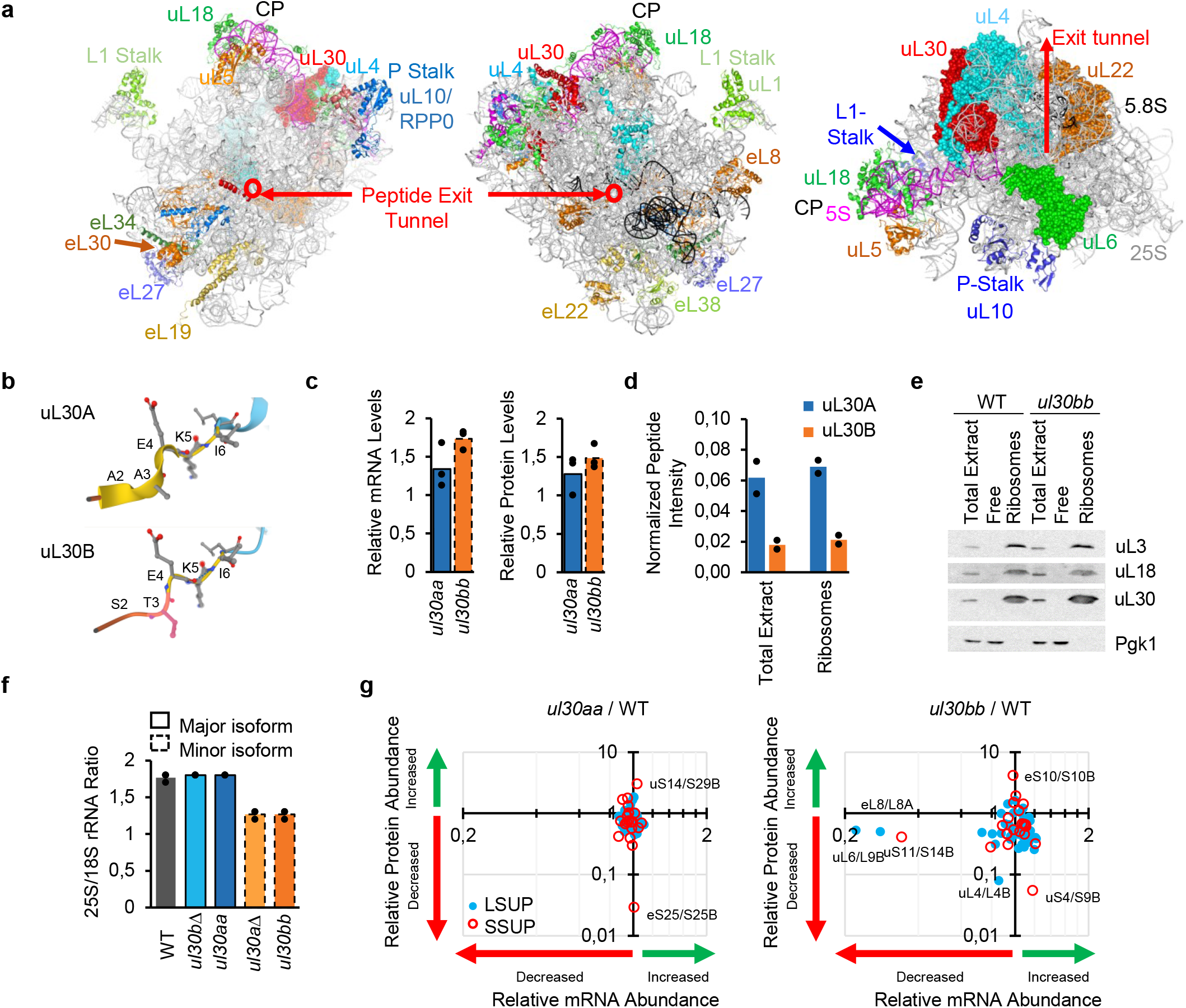
Impact of *uL30/RPL7* paralog on protein and ribosome production. **a,** Position of uL30/RPL7 protein within the 60S subunit. The ribosome structure was obtained from PDB file 5JUO originating from Ben-Shem et al., 2011, Science 334:1524. uL30 is indicated in red spheres. The rRNA and spatially or functionally related proteins are labeled. CP and P-stalk indicate the central protuberance and P-stalk base. **b,** A N-terminal ribbon structure is specific to uL30A as predicted by secondary structure comparison of the N-termini of uL30 paralogs from AlphaFold Protein Structure Database. **c,** Total uL30 mRNA was detected in *ul30aa* or *ul30bb* cells using common primers in qRT-PCR and shown relative to wild-type (left panel). The amount of uL30 protein in *ul30aa* or *ul30bb* cells was determined relative to wild-type after normalization to 60S RPs using Swath MRM (right panel). The data is from 3 biological replicates. **d,** Paralog-specific peptides were detected as described in (**c**) in total extract and ribosomes containing fractions from sucrose sedimentation in 2 biological replicates. **e,** Western blot of uL30, uL3, uL18 and the control Pgk1 proteins from total extracts and sucrose sedimentation fractions. **f,** The ratio of the 25S to 18S rRNA was determined using capillary electrophoresis. The data is from 3 biological replicates. **g,** Dot plot showing the changes in abundance of the mRNA and proteins produced by different RPGs in cells expressing one or the other paralog of *uL30/RPL7* using RNA-seq and Swath MRM and plotted relative to wild-type. LSU and SSU proteins are indicated by blue and red circles, respectively. The most affected RPGs in the homogenized strains are labeled. The RNA sequencing data are average of 2 biological replicates, the proteomic data are average of 3 biological replicates.

**Extended Data Fig. 3.**
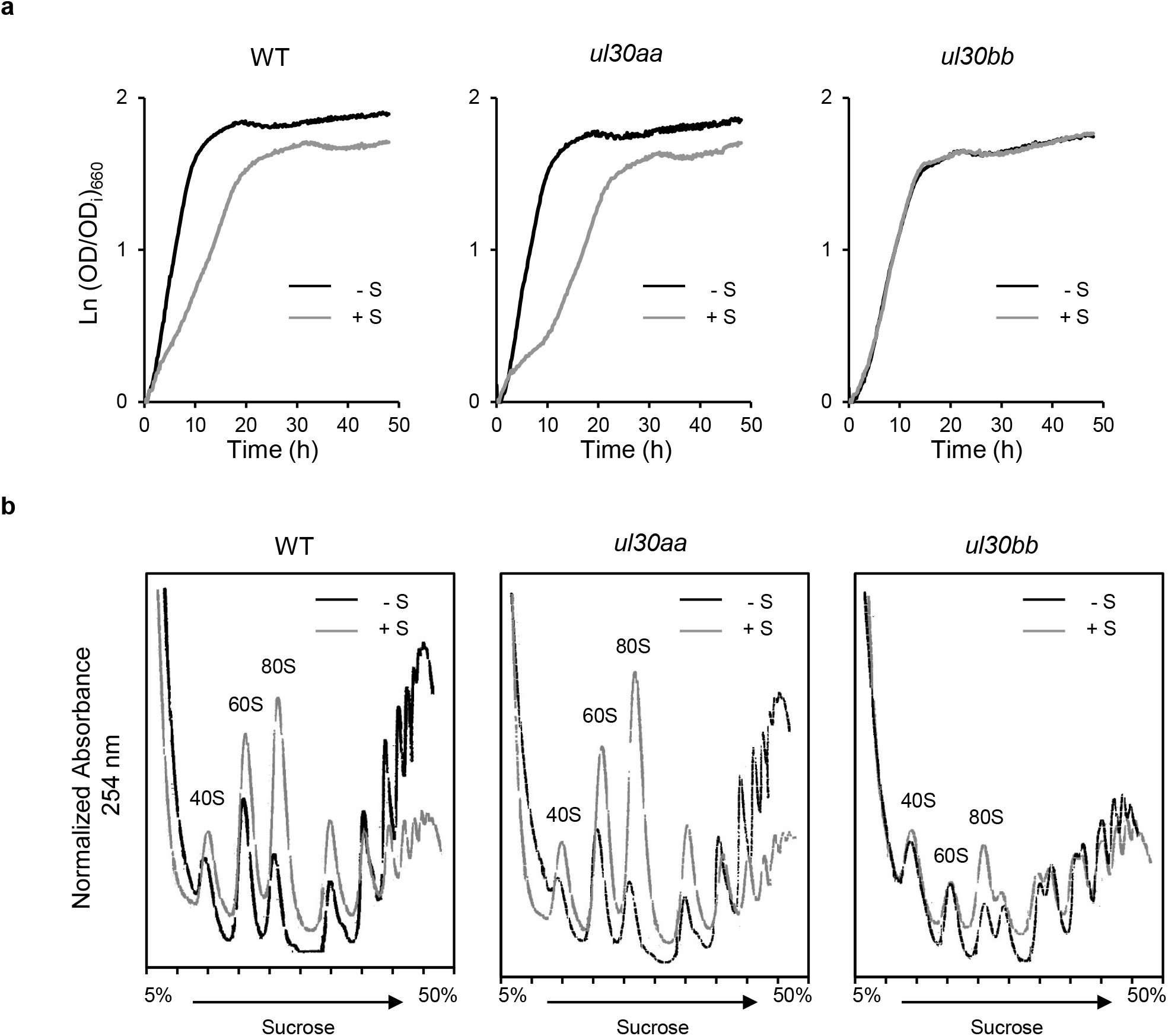
Features and impact of uL30/RPL7 paralogs on cell growth and translation. **a,** Comparison of the growth profile of wildtype (WT) and homogenized (*ul30aa* and *ul30bb*) strains in the presence (+ S) or the absence (- S) of staurosporine. **b,** Comparison of the polysome profile generated from wildtype (WT) and homogenized (*ul30aa* and *ul30bb*) strains strains grown in the presence (+ S) or the absence (- S) of staurosporine. The position of the monosomes and subunits is shown on top. Curves shown are representative examples of 3 biological repeats.

**Extended Data Fig. 4.**
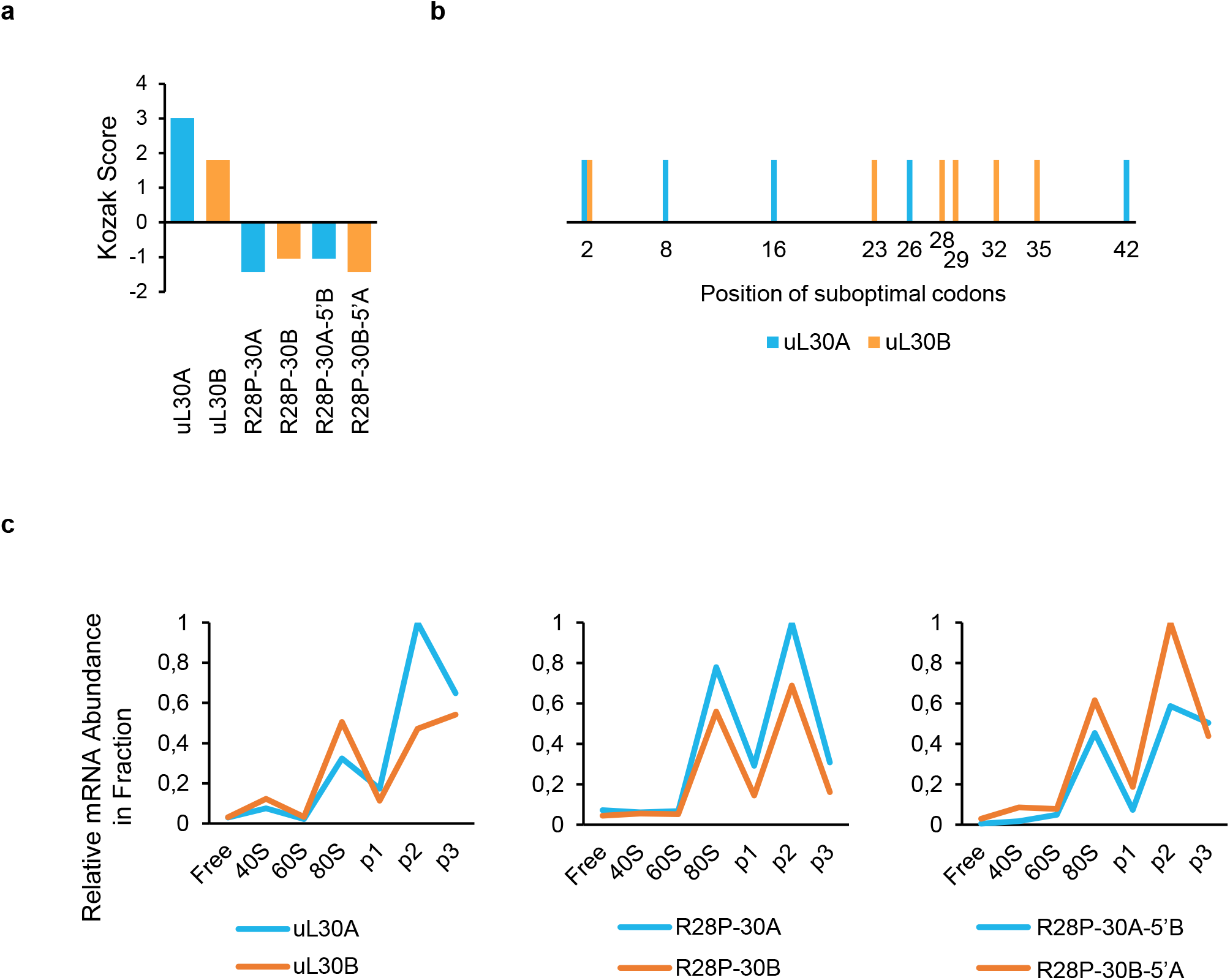
uL30A paralog mRNA is preferentially translated. **a,** The Kozak score (optimal start codon context) was calculated for each construct and presented in the form of a bar graph. **b,** Graph mapping the distribution of suboptimal codons in the first 42 amino acids of uL30A and uL30B. **c,** Translation profiles of uL30A and uL30B in WT and constructs. uL30A and uL30B mRNA from sucrose sedimentation fractions was determined using qRT-PCR. Enrichments were normalized to the highest point per strain pairs and curves represent the average of 2-3 biological replicates.

**Extended Data Fig. 5.**
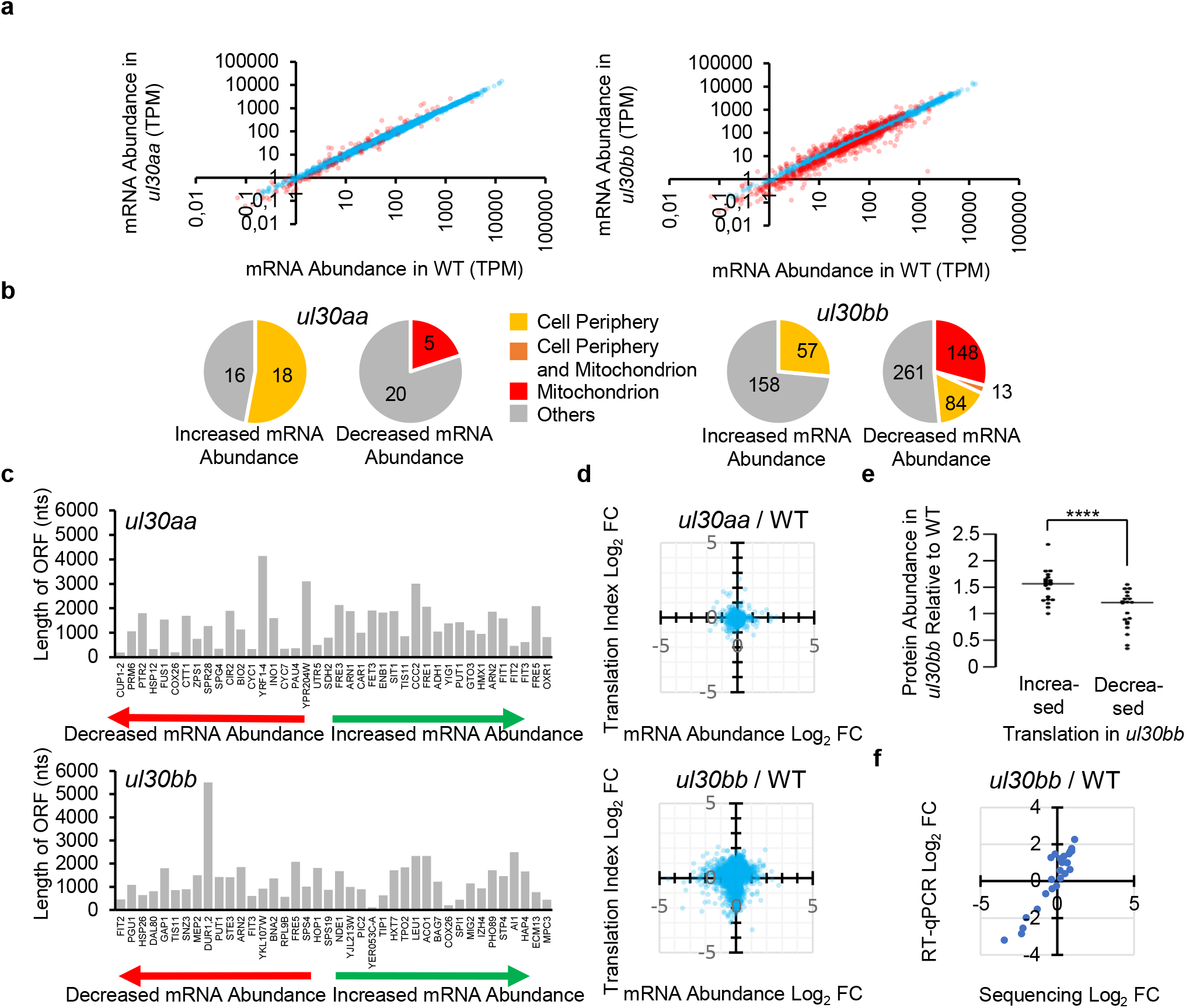
Impact of the homogenization of *uL30/RPL7* paralogs on RNA abundance. **a,** Comparison of expression levels from total RNA sequencing. Genes that vary relative to the wild type by >Log_2_ 0.5 are shown in red. Data points are an average of 2 biological repeats. **b,** Distribution of the number of genes present in enriched component gene ontology categories (p <0.001) for genes with changes in expression identified in (**a**). An overlap of 13 genes was found in the cell periphery and mitochondrion categories within the mRNAs under-expressed in *ul30bb*. **c,** The top 20 genes over- or under-expressed in *uL30aa* strain (top panel) or *ul30bb* (lower panel) are plotted relative to the length of their ORF in nucleotides. Genes are ordered on the X-axis as function of the magnitude of their change in expression. **d,** Comparison between the change in translation index and mRNA abundance in *ul30aa* and *ul30bb* relative to wild-type strain. The translation index was calculated described in Fig. 3a. **e,** Protein abundance of the top 20 genes with altered translation in *ul30bb* strain was determined using Swath Multiple Reaction Monitoring (MRM) and compared to WT after normalization to Pgk1p. The data points are average of 3 biological replicates, the line represent the median (**** p< 0.0001 by two-tailed t-test assuming unequal variance). **f,** Comparison of the translation index measured by RNA sequence and qRT-PCR. The translation index of 20 genes with altered translation in *ul30bb* strain was calculated using qRT-PCR and compared to that detected by RNA sequence. The two data set come from 2 biological repeats and exhibited Pearson correlation coefficient of 0.887.

**Extended Data Fig. 6.**
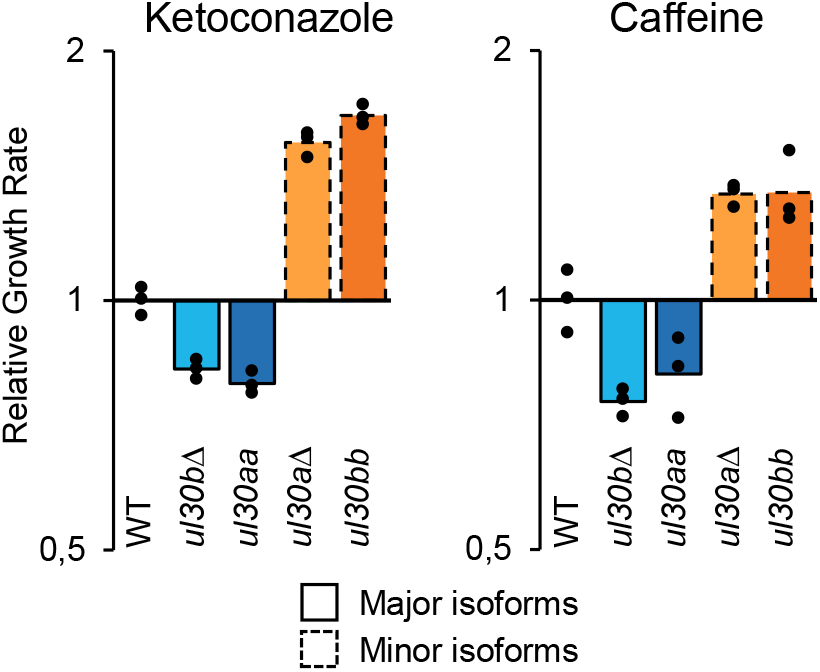
The minor paralog of *uL30/RPL7* increases resistance to drugs inducing cell wall stress. Wild type, *ul30bΔ*, *ul30aΔ*, *ul30aa* and *ul30bb* cells were grown in the presence or the absence of ketoconazole (32 µg/ml, left panel) or caffeine (10 mM, right panel) and the effect on growth rate on the different strains shown relative to that of wild type. The bars represent the average effect observed in three independent biological replicas shown as data points.

**Extended Data Fig. 7.**
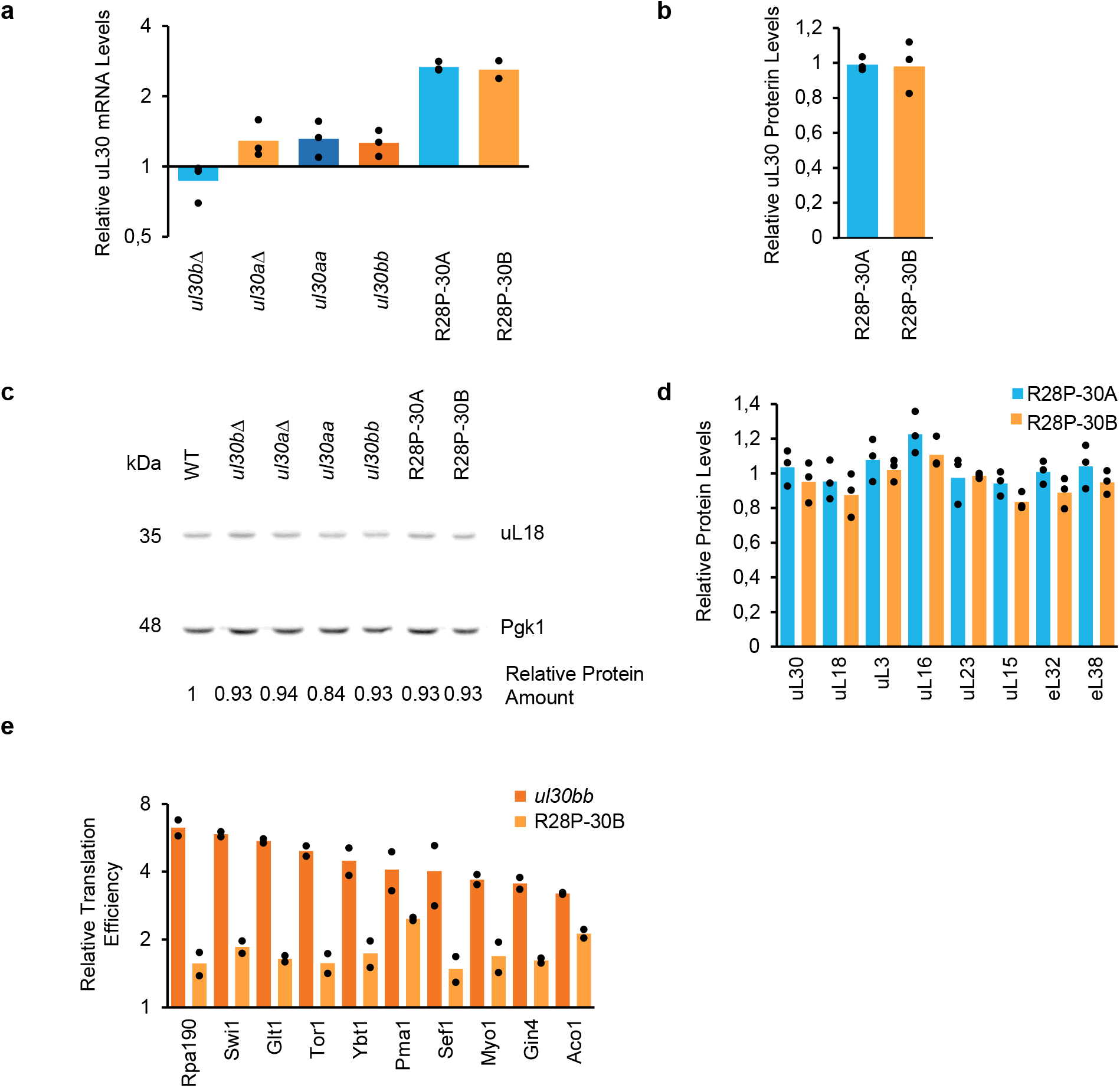
Comparison between RNA, protein and ribosome abundance generated from the chromosomal and plasmid-borne copies of *uL30/RPL7*. **a,** *uL30/RPL7* mRNA level from either plasmid or the chromosomal locus was determined using qRT-PCR and shown relative to the WT for 2-3 biological replicates. **b,** Quantification of uL30/RPL7 paralogs expressed from plasmids relative to WT after normalization to 60S RPs for 3 biological replicates using Swath MRM. Both strains express only one version of uL30. **c,** Western blot of uL18 with relative quantification to Pgk1 in strains expressing *uL30/RPL7* paralog from either the chromosomal locus or plasmid. **d,** Swath MRM was used to quantify 60S RPs normalized to Pgk1 in strains expressing *uL30/RPL7* from plasmids for 3 biological replicates. **e,** Translation index of mRNA in cells expressing uL30B from chromosomal copies (*ul30bb*) or from a plasmid (R28P-30B) relative to WT. The translation index of 10 long mRNAs (2337-7413 nts) were calculated using qRT-PCR as described in Figure 3a for 2 biological replicates.

**Extended Data Fig. 8.**
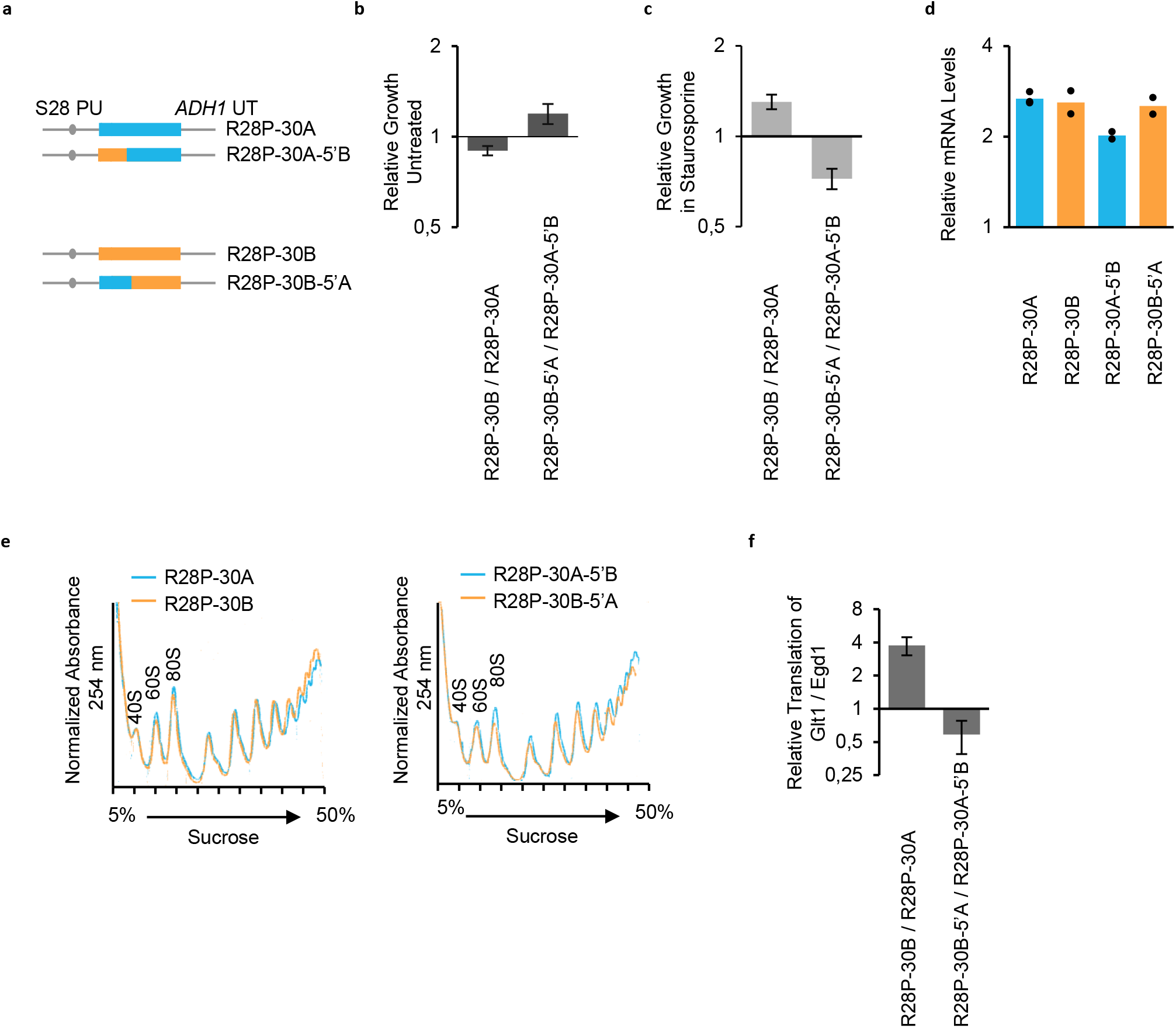
The difference in staurosporine resistance and mRNA size selection is included in the N-terminal part of uL30. **a,** The plasmid-borne uL30 paralogs and mutations used in **b-f** are schematically illustrated and the position of the mutations are indicated in the form of boxes. The ORFs of uL30A/RPL7A and uL30B/RPL7B are indicated as light blue and orange boxes. **b,** The relative growth rate of cells expressing one or the other uL30/RPL7 paralog from plasmid or the different mutations was determined in YC complete media and shown in the form of a bar graph with error bars representing the error on the ratio from the standard deviation of 3 biological replicates. **c,** The growth rate in the presence of staurosporine was calculated as in **b** for strains carrying the different constructs and relative growth of each plasmid pair indicated in the form of a bar graph for 2-4 biological Ns replicates. **d,** The mRNA produced from the different constructs was determined using RT-qPCR from 2-3 biological replicates. **e,** Comparison of the polyribosome profiles from cells expressing the different constructs. Representative curves from 3 biological replicates are shown. **f,** The ratio of mRNA with long (Glt1) and short (Egd1) open reading frame associated with heavy polyribosome was examined in cells expressing wild-type or mutated versions of uL30/RPL7 paralog using RT-qPCR as explained in Figure 3a. The error bars represent the error on the ratio from the standard deviation of 2-3 biological replicates.

